# Increased neurovirulence of omicron BA.5 and XBB variants over BA.1 in K18-hACE2 mice and human brain organoids

**DOI:** 10.1101/2022.12.22.521696

**Authors:** Romal Stewart, Kexin Yan, Sevannah A. Ellis, Cameron Bishop, Troy Dumenil, Bing Tang, Wilson Nguyen, Thibaut Larcher, Rhys Parry, Julian De Jun Sng, Alexander A. Khromykh, Robert K. P. Sullivan, Mary Lor, Frédéric A. Meunier, Daniel J. Rawle, Andreas Suhrbier

## Abstract

The reduced pathogenicity of the omicron BA.1 sub-lineage compared to earlier variants is well described, although whether such attenuation is retained for later variants like BA.5 and XBB remains controversial. We show that BA.5 and XBB isolates were significantly more pathogenic in K18-hACE2 mice than a BA.1 isolate, showing increased neuroinvasiveness, resulting in fulminant brain infection and mortality, similar to that seen for original ancestral isolates. BA.5 also infected human cortical brain organoids to a greater extent than the BA.1 and original ancestral isolates. In the brains of mice, neurons were the main target of infection, and in human organoids neuronal progenitor cells and immature neurons were infected. Although fulminant brain infection is not a feature of COVID-19, evidence for brain infection and brain damage in some COVID-19 patients with severe disease is becoming compelling, with the results herein suggesting that evolving omicron variants may have increasing intrinsic neuropathogenic potential.

## INTRODUCTION

The SARS-CoV-2 omicron lineage diverged considerably from earlier variants of concern, with the evolutionary origins remaining unclear, with the closest genetic common ancestor dating back to mid-2020 (Du et al., 2022; Mallapaty, 2022). Omicron viruses have spread faster globally than any previous variants; the BA.5 sub-lineage became the dominant SARS-CoV-2 virus in many countries (2022; Tanne, 2022), with XBB now spreading rapidly (Yue et al., 2023). Long-COVID is now well described for many variants of concern (Davis et al., 2023), and also occurs after infection with BA.5 (Qasmieh et al., 2023). Neurological and psychiatric manifestations represent a major component of COVID-19 and long-COVID (Davis et al., 2023; Islam et al., 2022; Monje and Iwasaki, 2022; Xu et al., 2022) and these remain for patients infected with omicron viruses (Chen et al., 2022; Cloete et al., 2022; Ludvigsson, 2022; Taquet et al., 2022), although extensive data specifically for BA.5 or XBB neurological manifestations have not yet emerged. The large number of changes in spike for omicron and omicron sub-lineages has rendered vaccination (Branche et al., 2022; Hansen et al.; Surie et al., 2022), many monoclonal antibody treatments (Takashita et al., 2022), and prior exposure with other variants (Suryawanshi et al., 2022) less protective.

There is increasing evidence for brain abnormalities in COVID-19 patients (Douaud et al., 2022; Graham et al., 2022; Karpiel et al., 2022; Ledford, 2022; Pelizzari et al., 2022; Sanabria-Diaz et al., 2022). Encephalitis is well documented, usually in hospitalized COVID-19 patients with severe disease, with encephalitis predisposing to poor outcomes and a higher risk of mortality (Altmayer et al., 2022; Chakraborty and Basu, 2022; Islam et al., 2022; Ong et al., 2023; Siow et al., 2021). The mechanism(s) whereby brain pathology/immunopathology and/or neuropathology might manifest in COVID-19 patients remains debatable, with the systemic cytokine storm and/or direct brain infection likely involved (Andrews et al., 2022; Aschman et al., 2022; Bauer et al., 2022a; Burks et al., 2021; Douaud et al., 2022; Fernandez-Castaneda et al., 2022; Rutkai et al., 2022; Samudyata et al., 2022; Silva et al., 2023; Zhang et al., 2020). A range of studies have now shown brain infection in COVID-19 patients (see Supplementary Table 1 for a full list and summary of findings). For instance, viral RNA or protein detected in the brains of 20-38% of patients that died of COVID-19 (Matschke et al., 2020; Serrano et al., 2022). A number of groups have also reported detection of viral RNA in cerebrospinal fluid of COVID-19 patients (Supplementary Table 1), including patients infected with omicron virus strains (Dang et al., 2022). Finally, COVID-19-associated damage to the brain is also likely to be associated with neurological manifestations of long-COVID (Davis et al., 2023; de Paula et al., 2023; Ferrucci et al., 2023; Rothstein, 2023), and such damage may also contribute to the continuing excess deaths arising from the COVID-19 pandemic (De Hert et al., 2021; Li et al., 2021; Msemburi et al., 2023).

The K18-hACE2 mouse model represents a model of severe COVID-19 that significantly recapitulates the lung pathology and inflammatory pathways seen in humans, and has been widely used for assessing new interventions and for studying SARS-CoV-2 biology (Bishop et al., 2022; Dong et al., 2022; Yinda et al., 2021). Infection of K18-hACE2 mice with original ancestral isolates via intranasal inoculation, usually results in fulminant and lethal brain infections, with virus likely entering the brain via the olfactory epithelium, across the cribriform plate and into the olfactory bulb (Carossino et al., 2022; Dumenil et al., 2022; Morgan et al., 2023; Olivarria Gema et al., 2022; Rothan et al., 2022; Vidal et al., 2022). This route of entry into the brain has been implicated in a non-human primate study (Jiao et al., 2021) and may also be relevant for COVID-19 patients, although the fulminant brain infection seen in these mice is not a feature of COVID-19 (Awogbindin et al., 2021; Beckman et al., 2022; Bulfamante et al., 2020; Matschke et al., 2020; Meinhardt et al., 2021; Serrano et al., 2022). Neurons represent a target of SARS-CoV-2 infection in brains of K18-hACE2 mice (Morgan et al., 2023; Rothan et al., 2022; Seehusen et al., 2022), non-human primates (Beckman et al., 2022), and hamsters (Ferren et al., 2021), with infection of neurons also observed in COVID-19 patients (Crunfli et al., 2022; Shen et al., 2022; Song et al., 2021). Reduced brain infection and the ensuing reduction in mortality after infection of K18-hACE2 mice with omicron BA.1 (Halfmann et al., 2022; Shuai et al., 2022; Tarres-Freixas et al., 2022) has been viewed as evidence that these viruses are intrinsically less pathogenic (Shrestha et al., 2022; Sigal, 2022).

*In vitro* organoid systems provide another method for evaluating intrinsic pathogenicity, with infection of brain organoid systems well described (Hou et al., 2022; Mesci et al., 2022; Song et al., 2021). Infection of neuronal cells usually results in their demise (Mesci et al., 2022), with type I IFN induction seen in some studies (Hou et al., 2022) but not others (Song et al., 2021).

Controversy surrounds the issue of whether BA.5 has altered intrinsic pathogenicity compared with earlier omicron variants. Some hamster studies have suggest increased lung infection/pathology for BA.5 (Kimura et al., 2022; Ong et al., 2023), whereas other studies using hamsters and/or K18-hACE2 mice argued that the reduced pathogenicity seen for the early omicron variants (Shuai et al., 2022) was retained by BA.5 (Rizvi et al., 2022; Uraki et al., 2022) and XBB (Tamura et al., 2023). Similarly, while some human studies report increased pathogenicity for BA.5 (Hansen et al., 2023; Kang et al., 2023; Kouamen et al., 2022; Russell et al., 2023), others report no significant changes (Davies et al., 2023; Wolter et al., 2022). XBB variants appear to have increased transmission potential (Islam et al., 2023), as well as enhanced receptor binding affinity, although the implications for pathogenicity remain to be established (Yue et al., 2023). Assessments of intrinsic pathogenicity of new variants in human populations are complicated by the overall rising levels of protective immunity due to vaccinations and/or past infections, which would tend to reduce the clinical severity for later COVID-19 waves (Sigal, 2022; Wolter et al., 2022). In contrast, the increased risk of mortality, hospitalization and acute and post-acute sequelae seen for repeated infections (Bowe et al., 2022), would tend to increase the clinical severity for later waves. Perhaps also pertinent for any such assessments is the diversifying pattern of COVID-19 and long-COVID disease manifestations (Kouamen et al., 2022), with involvement of non-pulmonary organs/systems increasingly being recognized (Davis et al., 2023; El-Kassas et al., 2023; Nchioua et al., 2022; Normandin et al., 2023).

Herein we illustrate and characterize the increased levels of brain infection and mortality for BA.5 and XBB versus BA.1 infected K18-hACE2 mice. We also show that BA.5 productively infected human cortical brain organoids significantly better than BA.1. These results argue that BA.5 and XBB may have increased intrinsic neuropathogenic potential over BA.1.

## RESULTS

### Omicron BA.5 and XBB are lethal in K18-hACE2 mice

Infection of K18-hACE2 mice with original ancestral isolates of SARS-CoV-2 is well described and results in weight loss and mortality by ≈ 5 days post infection (dpi) (Amarilla et al., 2021; Carossino et al., 2022; Dumenil et al., 2022; Kumari et al., 2021; Yu et al., 2022; Zheng et al., 2021). We re-illustrate this phenomena herein using an original ancestral isolate (SARS-CoV-2_QLD02_) and K18-hACE2 mice, with the ethically defined end-point of >20% weight loss reached by 4-5 dpi (Fig. 1A, Original). An omicron BA.1 isolate (SARS-CoV-2_QIMR01_) was substantially less virulent, with only 20% of mice showing weight loss >20% requiring euthanasia by 9/10 dpi (Fig. 1A, Omicron BA.1; Supplementary Fig. 1A). The reduced pathogenicity of BA.1 isolates in K18-hACE2 mice is consistent with previous reports (Chen et al., 2023; Halfmann et al., 2022; Shuai et al., 2022; Tarres-Freixas et al., 2022). In contrast, infection of K18-hACE2 mice with an omicron BA.5 isolate (SARS-CoV-2_QIMR03_) or an omicron XBB isolate (SARS-CoV-2_UQ01_) resulted in severe weight loss requiring euthanasia in nearly all mice by 4-7 dpi (Fig. 1A, BA.5, XBB).

**Figure. 1.**
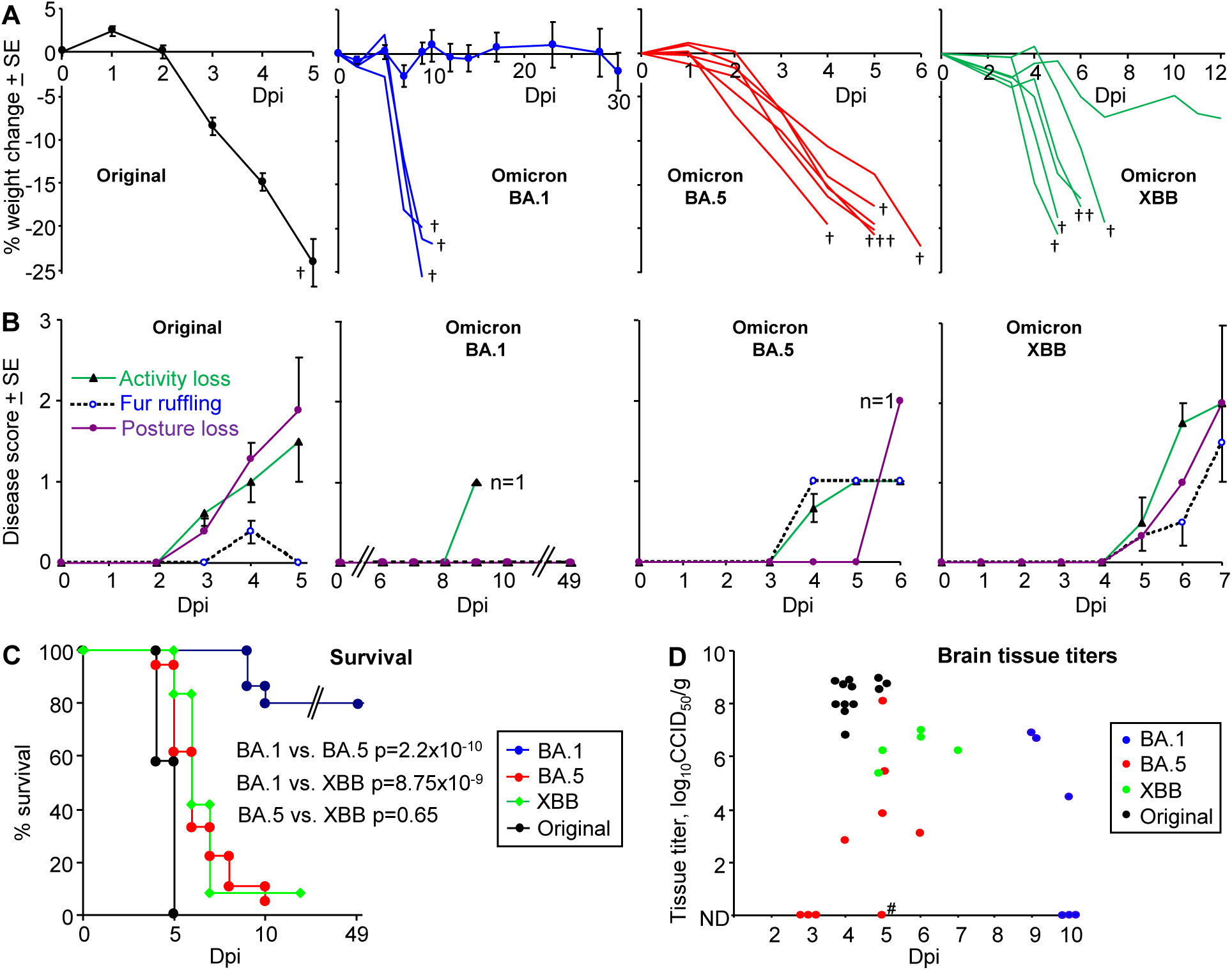
K18-hACE2 mice infected with an original ancestral isolate, omicron BA.1, BA.5 and XBB isolates. (A) Percent weight change post infection. † mice reached ethically defined end points for euthanasia. Original ancestral isolate (n=5-12 mice measured per time point), means and SEs are plotted for all mice (data from 3 independent experiments). BA.1, three mice showed weight loss >20% (requiring euthanasia, †) and are graphed individually; means and SE are plotted for the surviving mice (n=4-15 mice measured per time point). BA.5 (n=6) graphed individually. XBB (n=6) graphed individually. (B) Disease scores for the indicated overt disease symptoms for animals described in A. For BA.1 no overt symptoms were seen except in a single animal on 9 dpi, with this animal one of the 3 that required euthanasia due to weight loss. For BA.5 the remaining mouse (n=1) on 6 dpi showed a Posture score of 2. For XBB the one surviving mouse recovered after 7 dpi. (C) Kaplan Meyer plot showing percent survival; n=12 for original, n=12 for XBB, n=18 for BA.5 and n=24 for BA.1 (data from 2-3 independent experiments). Statistics by log-rank tests. (D) Viral tissue titers in brains. All mice with weight loss engendering euthanasia had detectable viral titers in brain (determined by CCID_50_ assays); the one exception (#) was IHC positive. Mice euthanized on 3 and 10 dpi with no detectable brain titers were symptom free. ND – not detected (limit of detection ≈ 2 log_10_CCID_50_/g). (Data from 2-3 independent experiments).

BA.5 and XBB infected mice also showed more overt disease symptoms than BA.1, with XBB showing disease score comparable with an original strain isolate (Fig. 1B). Consistent with previous reports (Bauer et al., 2022b; Seehusen et al., 2022), the majority of BA.1 infected mice showed no symptoms.

Kaplan Meier plots illustrate a highly significant difference between BA.1 vs. BA.5 and BA.1 vs. XBB, with no significant difference between BA.5 and XBB (Fig. 1C). Mortality from BA.5 and XBB was delayed when compared with the original ancestral isolate, although the mean delay was ≤ 2 days (Fig. 1C). The results for BA.5 contrast with a recent publication reporting that the reduced pathogenicity of early omicron sub lineages was retained for BA.5 (Uraki et al., 2022).

All mice that were euthanized due to reaching ethically defined end points (primarily <u>reaching</u> 20% weight loss), had detectable brain titers (Fig. 1D), with the exception of one BA.5-infected mouse whose brain was, however, subsequently found to be positive by IHC (Fig. 1D). Overall, such euthanized mice infected with the original ancestral isolate had >2 logs higher brain titers than euthanized mice infected with the omicron isolates (Fig. 1D). At euthanasia, there were no significant differences in brain titers amongst mice infected with the three omicron isolates (Fig. 1C). Six symptom-free mice had no detectable brain titers (Fig. 1D, ND, 3 and 10 dpi). Lung titres are shown in Supplementary Fig. 1B.

### Immunohistochemistry of BA.5 and XBB brain infection in K18-hACE2 mice

The fulminant brain infection seen after infection of K18-hACE2 mice with original ancestral isolates is well described, with widespread infection of neurons in various brain regions, including the cortex (Carossino et al., 2022; Morgan et al., 2023; Rothan et al., 2022; Seehusen et al., 2022; Vidal et al., 2022). A similar pattern of brain infection was observed using our K18-hACE2 mice and an original ancestral isolate, with immunohistochemistry (IHC) undertaken using a recently developed anti-spike monoclonal antibody (SCV2-1E8) (Morgan et al., 2023) (Supplementary Fig. 2A).

IHC staining of brains of K18-hACE2 mice infected with BA.5 or XBB also showed widespread infection of cells in the cortex, as well as the hippocampus and the hypothalamus (Fig. 2A,B,C). Viral RNA and protein has been detected in the cortex (Shen et al., 2022; Song et al., 2021) and hypothalamus (Stein et al., 2022) of post-mortem COVID-patients. Viral protein has also been identified in the hippocampus of such patients (Emmi et al., 2023), with disruption of the hippocampus also reported (Douaud et al., 2022; Radhakrishnan and Kandasamy, 2022). In the hippocampus of K18-hACE2, viral antigen could also be clearly seen in dendrites and axons (likely neural) (Fig. 2B,C, right hand panels), with viral antigen staining in neurites previously shown in human brain organoids (Bullen et al., 2020).

**Figure 2.**
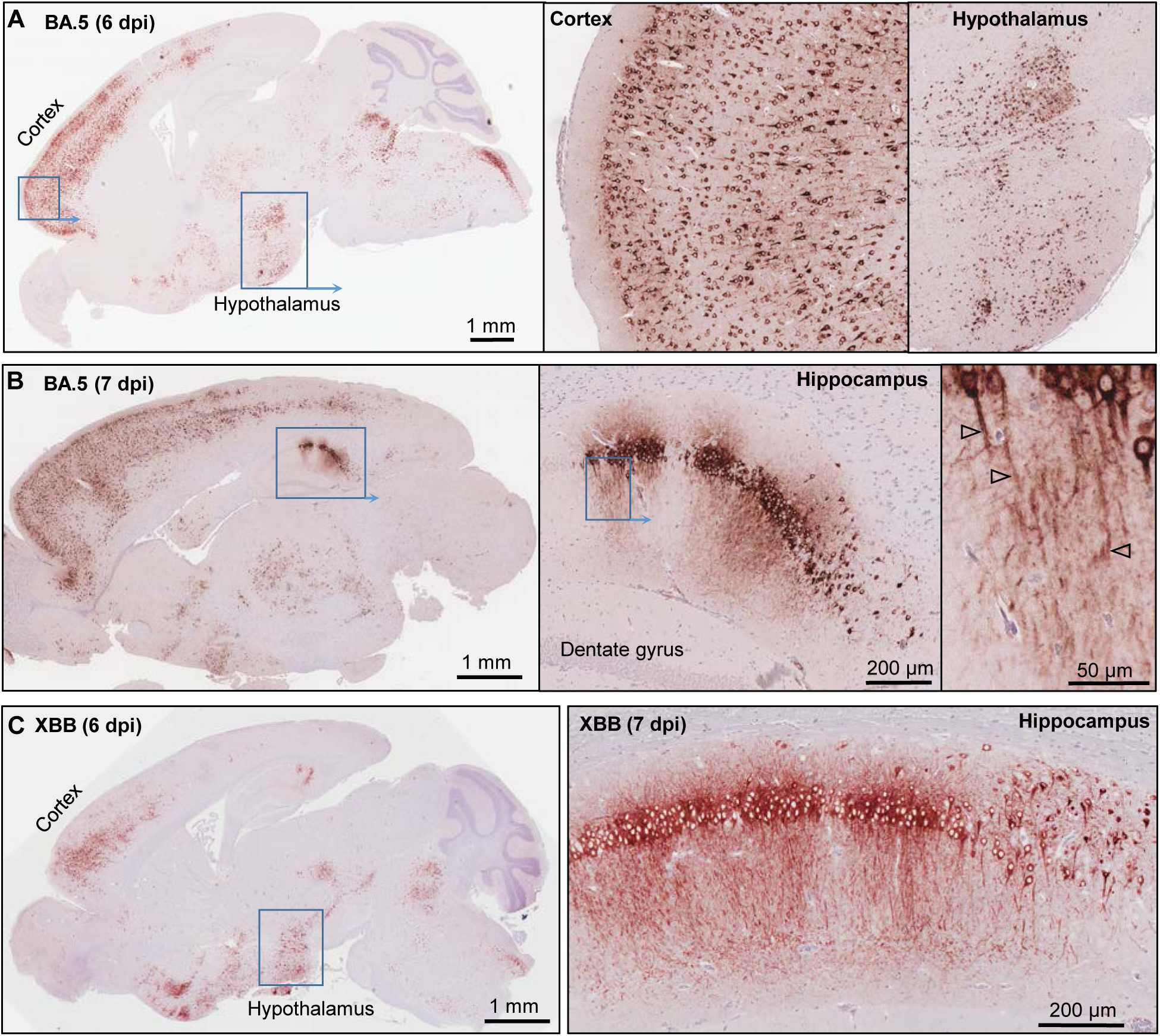
Immunohistochemistry of brains of BA.5 infected K18-hACE2 mice using an anti-spike monoclonal antibody. (A) Brain of BA.5-infected K18-hACE2 mouse 6 dpi showing IHC staining in the cortex and hypothalamus. Insert enlargements on the right. (B) As for A showing staining of the hippocampus 7 dpi. Far right shows staining of the axons (arrowheads). (C) XBB-infected K18-hACE2 mouse showing IHC staining in the cortex and hypothalamus, 6 dpi. Staining of the hippocampus for a mouse euthanized 7 dpi.

As described above, in the K18-hACE2 model, brain infection is generally associated with weight loss and mortality (Fumagalli et al., 2022) (with mice euthanized after reaching ethically defined end points). Perhaps of note, even low levels of IHC-detectable BA.5 infection was associated with weight loss that required euthanasia (Fig. 1D, Supplementary Fig. 2B), illustrating that such fatal outcomes do not require a fulminant brain infection.

### BA.5 infects neurons in K18-hACE2 mice

To confirm infection of neurons by BA.5, the cortex of infected K18-hACE2 mice were co-stained with the anti-spike monoclonal antibody and anti-NeuN, a neuronal nuclear antigen marker. Extensive co-localization within the same cells was observed (Fig. 3, Neurons) illustrating that neurons are a primary target of BA.5 infection in K18-hACE2 mouse brains. Co-staining with anti-spike and anti-Iba1, a pan-microglia marker, showed minimal overlap with anti-spike (Fig. 3, Microglia), arguing that microglia are not a major target of infection. Occasional overlap (yellow) may be due to phagocytosis of debris from virus-infected cells may arise from the recently described fusion events (Martínez-Mármol et al., 2023). Despite being surrounded by infected neurons, most microglia retaining their ramified morphology, although some cells with bushy and amoeboid morphology were present (Fig. 3, Microglia) indicating activation-associated retraction of processes (Pinto and Fernandes, 2020). Microglia activation was also indicated by histology and RNA-Seq (see below). Although occasionally seen (Supplementary Fig. 3A), anti-GFAP staining was minimal around infected neurons (Fig. 3, Reactive astrocytes), arguing that astrocytes are largely not being activated. RNA-Seq also did not identify GFAP as a significantly up-regulated gene, nor did bioinformatics identify an astrocyte activation signature (see below; Supplementary Table 2).

**Figure 3.**
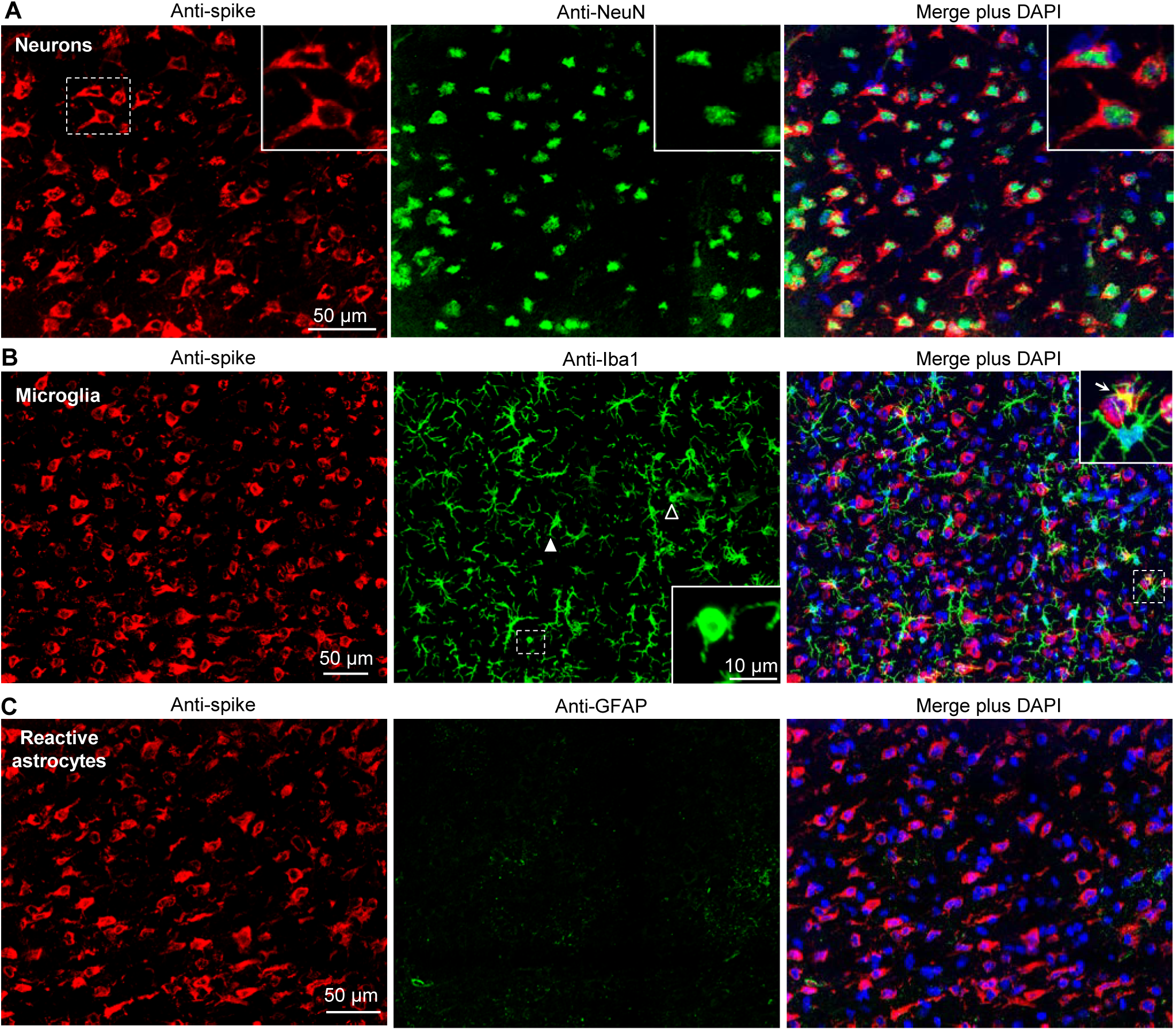
Dual labelling fluorescence immunohistochemistry of BA.5 infected brains from K18-hACE2 mice. (A) Sections from formalin fixed and paraffin embedded brains from BA.5 infected K18-hACE2 mouse brains (5 dpi) were analyzed by immunofluorescence. Sections were stained by indirect immunofluorescence with the anti-spike monoclonal antibody (red), and antibodies specific for a neuronal nuclear marker - NeuN (green). Dashed white box (left) indicate cells enlarged in the three inserts (top right). (B) As for A but costaining with the pan-microglial marker - Iba1 (green). Anti-Iba1 (middle panel); an enlargement of the dashed box (middle panel) is shown in the insert (bottom right), and indicates a microglial cell with amoeboid morphology. Another microglial cell with amoeboid morphology is indicated with a white unfilled arrowhead. White filled arrowhead indicates a microglial cell with bushy morphology. Merged plus DAPI (right panel); an enlargement of the dashed box is shown top right, with arrow indicating yellow (red green overlap), possibly associated with phagocytosis of infected cell debris. (C) As for A but costaining with a reactive astrocyte maker - GFAP (green).

### Brain lesions identified by H&E in BA.5 and XBB infected K18-hACE2 mice

The brains of BA.5 infected K18-hACE2 mice showed a number of lesions that have also been observed in COVID-19 patients and/or primate models. Neuron vacuolation (hydropic degeneration) was clearly evident (Fig. 4A), and has been reported previously for infection of K18-hACE2 mice with an original ancestral isolate (Vidal et al., 2022), and was also observed in non-human primate (NHP) model of SARS-CoV-2 infection (Rutkai et al., 2022). The presence of viral antigen in the cortex was associated with apoptosis (Supplementary Fig. 3B) and a high intensity of H&E-detectable lesions (primarily vacuolation), but was not associated with local immune cell infiltrates (Supplementary Fig. 4). The lack of infiltrates around areas of infection has also been noted in COVID-19 patients (Song et al., 2021). Perivascular cuffing (Fig. 4B) is well described in histological examinations of brains from deceased COVID-19 patients (Awogbindin et al., 2021; Matschke et al., 2020; Rosu et al., 2022; Schwabenland et al., 2021; Serrano et al., 2022). Other lesions observed in the BA.5-infected K18-hACE2 mouse brains, that have also been described in post-mortem COVID-19 patients, include perivascular oedema (Fig. 4C) (Maiese et al., 2021; Martin et al., 2022; Pajo et al., 2021), occasional microglial nodules (Fig. 4D) (Al-Dalahmah et al., 2020; Awogbindin et al., 2021; Matschke et al., 2020; Schwabenland et al., 2021), and occasional small hemorrhagic lesions (Fig. 4E) (Mukerji and Solomon, 2021; Rosu et al., 2022). XBB infected mouse brains showed very similar lesions (Fig. 4F, microgliosis, perivascular cuffing, perivascular oedema). Chromatolysis, indicative of injury, was also evident in hippocampal neurons (Fig. 4G, dashed box) in the IHC positive region identified in Fig. 2C.

**Figure 4.**
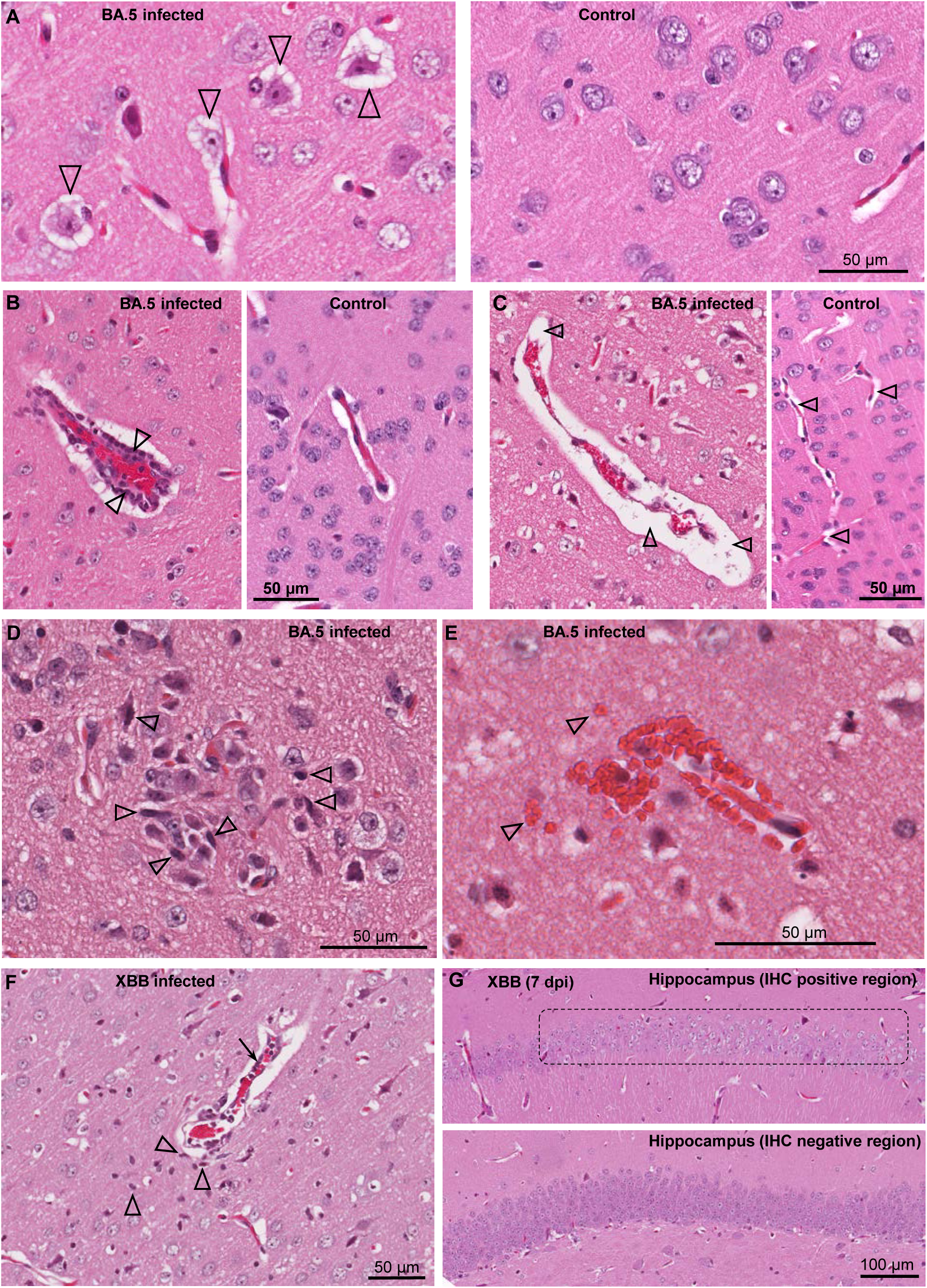
Histological lesions in brains of BA.5 and XBB infected K18-hACE2 mice. H&E staining of brains from BA.5 (A-E) and XBB (F-G) infected K18-hACE2 mice. (A) Neuron vacuolation (hydropic degeneration) of neurons (arrowheads) in the cortex 4 dpi. A control brain from mice inoculated with UV-inactivated BA.5 is shown on the right. (B) Perivascular cuffing. A venule (red blood cells in the center) is surrounded by leukocytes (two leukocytes are indicated by arrowheads), 7 dpi. A control venule is shown on the right. (C) Focal vasogenic edema; fluid filled perivascular space (arrowheads), 6 dpi. A control is shown on the right, with arrowheads showing normal perivascular spaces. (D) Microglial nodule; accumulation of microgliocytes (some typical microgliocytes indicated with arrowheads), 6 dpi. (E) Small hemorrhagic lesion, 7 dpi (arrowheads indicate some extravascular red blood cells). (F) Lesion in the cortex (from the IHC positive region in Fig. 2C, left panel) showing perivascular cuffing (arrow), microgliosis (arrowheads) and vasogenic edema (as in C), 6 dpi. (G) Loss of haematoxylin staining of neurons in the hippocampus (chromatolysis) in the anti-spike positive region shown by IHC in Fig. 2C (right panel). An IHC spike negative region from the hippocampus of the same mouse is shown as a control (bottom image).

### RNA-Seq of BA.5-infected K18-hACE2 mouse brains

Mice were infected as in Fig. 1 (BA.5) and euthanized when weight loss reached the ethically defined endpoint of ≈ 20% (Supplementary Fig. 5A). Control mice received the same inoculation of UV-inactivated BA.5. Brains were examined by RNA-Seq (BioProject ID: PRJNA911424), with the PCA plot shown in Supplementary Fig. 5B and viral RNA levels in Supplementary Fig. 5C. Differentially expressed genes (DEGs) (q<0.05, n=437) were analysed by Ingenuity Pathway Analysis (IPA) (Bishop et al., 2022; Dumenil et al., 2022) (Supplementary Table 2). Selected representative IPA annotations, grouped by themes, are shown in Fig. 5A. The dominant annotations illustrate a cytokine storm, with the top cytokine Up Stream Regulators (USRs) including interferons both type II (IFNγ) and type I (IFNα2, IFNλ1 IFNβ1), as well as TNF, IL-1 and IL-6, all previously well described for SARS-CoV-2 infections (Bishop et al., 2022). The concordance for cytokine USRs for brain and lung infection, and for the three virus isolates, was high (Supplementary Fig. 5D), arguing that inflammatory responses are generally very similar for brain and lungs and for the different SARS-CoV-2 variants.

**Figure 5.**
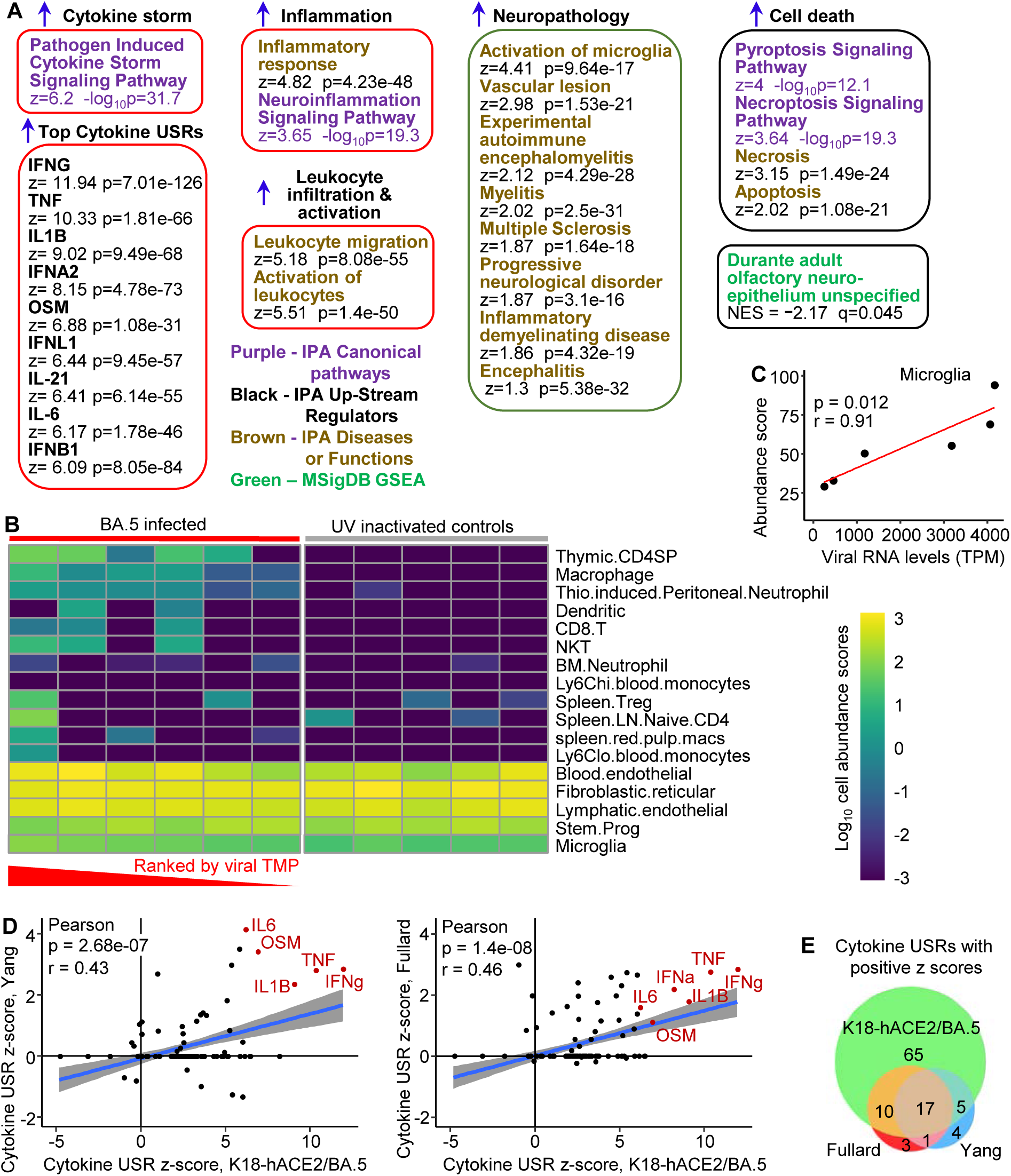
Transcriptome signatures in brains of K18-hACE2 mice infected with BA.5. (A) RNA-Seq of BA.5 infected brains (n=6) compared with brains of mice inoculated with UV-inactivated BA.5 (n=5) identified 437 DEGs. The DEGs were analyzed by Ingenuity Pathway Analysis (IPA) and GSEAs using the Molecular Signatures Data Base (MSigDB), with a representative sample of annotations shown and grouped by theme (a full list is provided in Supplementary Table 1). (B) The RNA-Seq expression data from brains of BA.5 infected K18-hACE2 mice were analyzed by SpatialDecon to provide estimates of cell type abundances. The BA.5 infected samples were ranked by viral RNA levels (highest to lowest). Cell types were clustered using the complete-linkage of Euclidean distance. (C) Relative cell type abundances for Microglia correlates with viral RNA levels. Statistics by Pearson correlation. (D) IPA cytokine USRs obtained from brains of BA.5 infected K18-hACE2 mice were compared with IPA cytokine USRs obtained using two DEG lists generated from publically available single-cell RNA-Seq data of selected brain tissues from deceased COVID-19 patients. Where a cytokine USR is identified in human but not mouse (or *vice-versa*), a z-score of zero is given to the latter. (E) Venn diagram showing overlaps between the cytokine USRs from the two human (Fullard and Yang) and the BA.5 infected K18-hACE2 mouse study (RNA-Seq data of brain tissues).

A large series of annotations were associated with leukocyte migration and activation (Supplementary Table 2), with the top two overarching annotations shown (Fig. 5A, Leukocyte migration, Activation of leukocytes). These annotations are consistent with the perivascular cuffing seen by H&E (Fig. 4B). An additional series of neuropathology-associated annotations were also identified with high z-scores and significance (Fig. 5A, Neuropathology). Activation of microglia and vascular lesions (Fig. 5A) were consistent with the histological findings (Fig. 4B,C,D). Multiple sclerosis-like features, myelitis, demyelination (Bellucci et al., 2021; Ismail and Salama, 2022) and encephalitis (Altmayer et al., 2022; Chakraborty and Basu, 2022; Islam et al., 2022; Siow et al., 2021) are also features described for COVID-19 patients. Apoptosis of neurons was reported in the NHP model (Rutkai et al., 2022), with pyroptosis in the CNS of COVID-19 patients also proposed (Sepehrinezhad et al., 2021). Gene Set Enrichment Analyses (GSEAs) using gene sets provided in MSigDB (≈ 50,000 gene sets) and in Blood Transcription Modules, generated broadly comparable results to those obtained from IPA (Supplementary Table 2). In addition, a significant negative enrichment (negative NES) for olfactory neuroepithelium genes (MSigDB) (Durante et al., 2020) was also identified (Fig. 5A), suggesting loss of cells in this tissue in BA.5-infected mouse brains. COVID-19-associated anosmia (loss of smell) in humans is likely associated with infection of the olfactory epithelium (Dumenil et al., 2022).

To provide insights into the nature of the leukocyte infiltrates, cell type abundance estimates were obtained from the RNA-Seq expression data using SpatialDecon (Danaher et al., 2022) (Fig. 5B). The inflammatory infiltrate appeared primarily to comprise immature CD4 T cells (Hosseinzadeh and Goldschneider, 1993), macrophages, neutrophils, dendritic cells, CD8 T cells and NKT cells, with increased cell abundance scores seen with increasing viral RNA levels (Fig. 5B, TPM - transcripts per million; Supplementary Fig. 5E). Although not substantial, increased cell abundance scores also increased with viral RNA levels for microglia (Fig. 5C).

In summary, the bioinformatic analyses illustrate that the inflammatory responses in BA.5-infected K18-hACE2 mouse brains are largely innate (4-6 dpi) and typical of acute SARS-CoV-2 infections, with many annotations consistent with histological findings and a series of studies in COVID-19 patients.

### Brain gene expression patterns in COVID-19 patients are recapitulated in SARS-CoV-2 infected K18-hACE2 mice

We previously illustrated that inflammatory pathways identified by RNA-Seq of lungs from COVID-19 patients showed highly significant concordances with SARS-CoV-2 infected lungs from K18-hACE2 mice (Bishop et al., 2022). Two publically available single-cell RNA-Seq data sets are available for selected human brain tissues (choroid plexus, medulla oblongata, and pre-frontal cortex) from deceased COVID-19 patients (Fullard et al., 2021; Yang et al., 2021). DEG sets from each tissue and cell-type were concatenated to create one overall DEG list for each of the two human studies. These DEG lists were analysed by IPA as above, and the cytokine USR z-scores compared with those obtained from brains of BA.5-infected K18-hACE2 mice. Highly significant concordances emerged for both human studies (Fig. 5D), with 80% of the cytokine USRs identified in humans also identified in K18-hACE2 mice. Thus, although fulminant lethal brain infection is not a feature of COVID-19 (Supplementary Table 1), the inflammatory signatures in brains of severe COVID-19 cases show a high degree of concordance with those seen in brains of K18-hACE2 mice.

Gene Set Enrichment Analyses (GSEAs) also illustrated that DEGs up-regulated in brains of COVID-19 patients (log_2_ fold change >1) (Fullard et al., 2021; Yang et al., 2021), were significantly enriched in the ranked gene list from brains of BA.5-infected K18-hACE2 mice (Supplementary Fig. 6). Thus patterns of gene expression in brain tissues from severe COVID-19 patients also showed significant overlap with those seen in BA.5-infected K18-hACE2 mice.

### Infection of human cortical brain organoids

Human, induced pluripotent cells (hiPSCs), derived from a primary dermal fibroblast line (HDFa) from a normal human adult, were used to generate approximately spherical, ≈ 2-3 mm diameter, “mini-brains” using a rotating incubator (Supplementary Fig. 7A). RNA-Seq and IHC illustrated that 30 day old organoids were comprised primarily of neural progenitor cells (expressing SOX2 and nestin) and immature neurons expressing MAP2 (Microtubule-Associated Protein 2) and TUBB3 (tubulin beta 3) (Supplementary Fig. 7B,C). Such organoids were infected with the BA.5, BA.1 and the original ancestral isolate (MOI≈ 1) and were cultured for 4 days. Dual labelling fluorescent IHC illustrated that BA.5 infected MAP-2-negative cells, and some MAP2-positive cells (Supplementary Fig. 8). The BA.5 virus infected substantially more cells in the organoids than the original ancestral (Fig. 6A) or the BA.1 viruses (Supplementary Fig. 9A). XBB also infected slightly more cells than BA.1 (Supplementary Fig. 9B). The small area infected with the original ancestral isolate (Fig. 6A, Original, insert) corresponded to an area of the organoid with IHC-detectable anti-hACE2 staining (Supplementary Fig. 9C). The overall expression of hACE2 mRNA was low, with TMPRESS2 mRNA often undetectable (Supplementary Fig. 9D). Viral titers in the supernatants of the organoid cultures increased over the 4 day period, with BA.5 titers significantly higher than BA.1 titers by 1-2 logs (Fig. 6B, p=0.007, 2, 3 and 4 dpi). XBB titers were also up to ≈1 log higher, which reached significance if data from 4 and 5 dpi were combined (Fig. 6B, p=0.009, 4 & 5 dpi). RNA-Seq of organoids harvested 4 dpi also illustrated that viral RNA levels were ≈ 25 fold higher for organoids infected with BA.5 than those infected with an original ancestral isolate (Fig. 6C).

**Figure 6.**
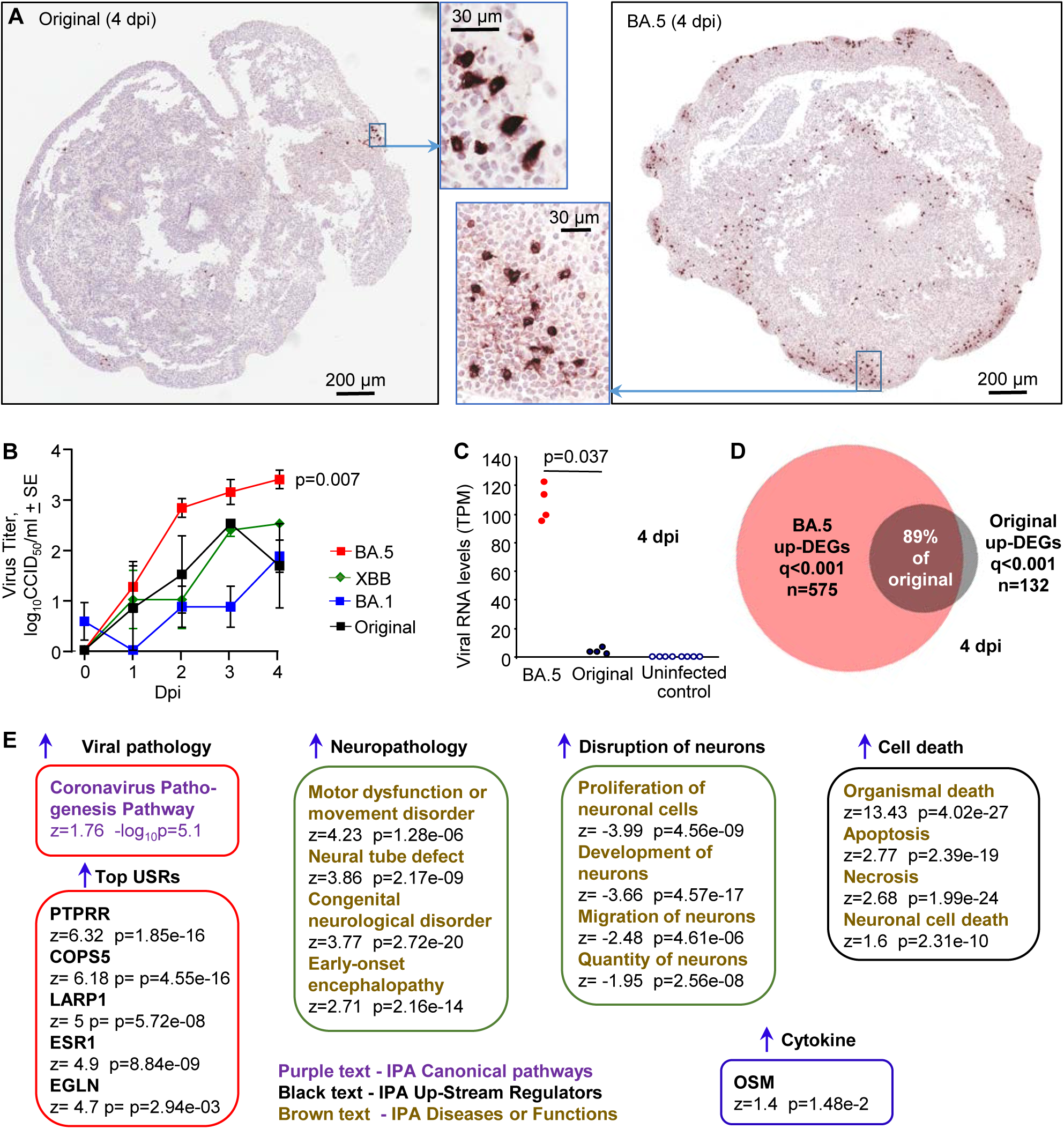
Infection of cortical brain organoids with original ancestral, BA. 1, BA.5 and XBB. (A) IHC of brain organoids 4 dpi with original ancestral and BA.5 using an anti-spike monoclonal antibody. IHC staining for XBB infected organoids is shown in Supplementary Fig. 9B. (B) Viral titers in the supernatant of the organoid cultures sampled at the indicated dpi. Data is derived from two independent experiments. Titers for BA.5 (n=8) vs. BA.1 (n=7) were significantly different (p=0.007) on days 2, 3 and 4. Titers for XBB (n=4) vs. BA.1 (n=7) were significantly different (p=0.009) when the data for 3 and 4 dpi were combined. Statistics by Kolmogorov-Smirnov tests. (C) RNA-Seq-derived viral reads counts for infected organoids, 4 dpi. (D) Venn diagram showing overlap of up-regulated DEGs for organoids infected with BA.5 and original ancestral. (E) DEGs (n=2390, q<0.001) generated from RNA-Seq of BA.5 infected organoids (n=4) 4 dpi vs. uninfected organoids (n=4) were analyzed by IPA. Selected and representative annotations are shown (for full data set see Supplementary Table 3).

RNA-Seq of BA.5 infected human cortical brain organoids (4 dpi) compared with uninfected organoids provided 2390 DEGs (q<0.001), of which 575 were up-regulated genes (Supplementary Table 3). RNA-Seq of original ancestral isolate-infected organoids provided 252 DEGs (q<0.001), of which 132 were up-regulated (Supplementary Table 3). Given the higher level of infection, more DEGs might be expected for BA.5. Of the 132 up-regulated DEGs, 118 were also identified in the BA.5 infected organoids (Fig. 6D), arguing that the original ancestral isolate is not inducing fundamentally different response in these organoids.

The 2390 DEGs for BA.5 were analysed by IPA (Supplementary Table 3), with “Coronavirus Pathogenesis Pathway” identified as a top canonical pathway (Fig. 6E). The top USRs were (i) PTPRR (a protein tyrosine phosphatase receptor), which was recently identified in a study of brains from SARS-CoV-2 infected hamsters and is associated with depression in humans (Serafini et al., 2022), (ii) COPS5 (COP9 signalosome subunit 5), whose mRNA is bound by SARS-CoV-2 NSP9, perhaps resulting in suppression of host responses (Banerjee et al., 2020), (iii) LARP1, a translational repressor that binds SARS-CoV-2 RNA (Schmidt et al., 2021), (iv) ESR1 (nuclear estrogen receptor), which is important for ACE2 expression (Oner et al., 2022), (v) EGLIN, oxygen sensors that target HIF α subunits for degradation, with HIF-1α promoting SARS-CoV-2 infection and inflammation (Tian et al., 2021). IPA Diseases and Functions feature identified a series of neuropathology-associated annotations, including a motor dysfunction signature, with motor deficits documented for severe COVID-19 patients (Graham et al., 2022). Consistent with the IHC data, a series of signatures describe disruption and death of neurons (Fig. 6E). No significant up-regulation of classical inflammation or IFN signatures were identified, with the possible exception of oncostatin M (OSM) (Fig. 6E). Serum concentrations of this IL-6 family pleiotropic cytokine show a strong positive correlation with COVID-19 severity (Arunachalam et al., 2020). However, OSM can also be secreted by neural cells, but in the brain it is thought often to play a neuroprotective role (Houben et al., 2019).

## DISCUSSION

We illustrate herein that BA.5 and XBB show greater propensities to enter the brain and infect neurons in K18-hACE2 mice when compared to BA.1. In addition, BA.5 showed an increased capacity to infect human brain organoids. Taken together these results argue that these two more recent omicron variants of concern may have enhanced intrinsic neuroinvasive and neurovirulence properties when compared to an earlier omicron variant. Some recent evidence from human studies may support this contention, with a higher proportion of patients infected with BA.5 developing anosmia, when compared with BA.1 (Hansen et al., 2023). Anosmia likely arises from infection of the olfactory epithelium, which contains a number of cell types including sustentacular cells, olfactory receptor neurons and basal cells. Infection of the olfactory epithelium also likely provides access to the brain (Dumenil et al., 2022; Fodoulian et al., 2020; Jiao et al., 2021). Anosmia was recently closely linked to long-lasting cognitive problems in COVID-19 patients (RADC, 2022). Thus, even if respiratory disease is unchanged or reduced, BA.5 may show increased risk of acute and long-term neurological complications over earlier omicron variants (Okrzeja et al., 2023).

The increased neuroinvasiveness of BA.5 and XBB over BA.1 in K18-hACE2 mice (Fig. 1A) may be associated with their enhanced fusion activity (Tamura et al., 2023; Tang et al., 2023), which is usually associated with enhanced ability to utilize TMPRSS2 and/or increased binding affinity for ACE2 (Aggarwal et al., 2022; Tamura et al., 2023). TMPRSS2 utilization is associated with virulence (Abbasi et al., 2021), even for omicron variants (Iwata-Yoshikawa et al., 2022), and is involved in neurotropism in K18-hACE2 mice (Li et al., 2021). An increased ability to infect, not just the TMPRSS2-positive sustentacular cells, but also TMPRSS2-low cells in the murine olfactory epithelium (Fodoulian et al., 2020), may thus promote entry of BA.5 and XBB into the brains of K18-hACE2 mice when compared with BA.1. In contrast to early omicron variants (Meng et al., 2022; Qu et al., 2023; Zhao et al., 2022), original ancestral isolates show a preference for TMPRSS2 utilization, and rapid fulminant infection of K18-hACE2 brains by such viruses is well described (Rothan et al., 2022; Seehusen et al., 2022). Interestingly, viral sequences from the brains of K18-hACE2 mice and hamsters infected with original ancestral isolates show loss of functional furin cleavage sites (Supplementary Fig. 10). TMPRSS2 mRNA expression levels in K18-hACE2 mouse brains are very low (Supplementary Table 2), so TMPRSS2-independent infection (Qu et al., 2023) would likely be selected as the virus spreads within the brains. Such furin cleavage site deletions were not seen in BA.1 or BA.5 sequences from brains of K18-hACE2 mice, likely because omicron viruses can already effectively use the endosomal pathway (Meng et al., 2022; Qu et al., 2023; Zhao et al., 2022). In summary, more efficient use of TMPRSS2-dependent infection by BA.5 and XBB (and original ancestral isolates) compared to BA.1, may promote entry into the brain of K18-hACE2 mice via the olfactory epithelium. Once in the brain, utilization of the endosomal pathway by omicron viruses (and original ancestral isolates with non-functional furin cleavage sites) allows a spreading infection in TMPRSS2-low brain cells (Fig. 1D, Brains).

Infection of brain organoids represents a measure of neurovirulence, rather than neuroinvasiveness, as access to cells in this *in vitro* system clearly does not require transit across the cribriform plate (Dumenil et al., 2022; Jiao et al., 2021). TMPRSS2 mRNA expression was even lower in the organoids (Supplementary Fig. 9C) than in K18-hACE2 brains, with infection of human neurons shown to be TMPRSS2-independent (Kettunen et al., 2023). This is consistent with the poor infection of organoids by the original ancestral isolate (Fig. 6A-C). BA.5 would thus appear to have an increased capacity for infection of brain organoids via a TMPRSS2-independent mechanism. This is not due to acquisition of hACE-2-independent infection capabilities (Yan et al., 2022) (Supplementary Fig. 11). Nor is this likely due to an increased ability of BA.5 to counter type I IFN activities (Guo et al., 2022), as such responses were not detected (Fig. 6E). hACE2 expression is low in the organoids, suggesting an increased affinity for hACE2 might promote BA.5 infection, with barely detectable levels of hACE2 able to support infection (Rawle et al., 2021). There are a number of differences in the spike protein between the BA.1 and BA.5 isolates including 9 amino acid changes in the receptor binding domain (Supplementary Fig. 12), with BA.5 affinity for hACE2 reported as slightly higher in two studies (Tuekprakhon et al., 2022; Wang et al., 2022), but unchanged in a third (Cao et al., 2022). More efficient use by BA.5/XBB of co-receptors such as neuropilin (Cantuti-Castelvetri et al., 2020; Kong et al., 2022) or heparin sulphate proteoglycans (Guimond et al., 2022) might also be involved. Non-spike changes may also play a role (Chen et al., 2023).

The evolution of SARS-CoV-2 shows a strong tendency towards increased transmission and immune evasion. Both are achieved by more rapid and fulminant infection of the upper respiratory tract (Carabelli et al., 2023), which in turn may increase the opportunities for virus to infect the brain (Dumenil et al., 2022; Jiao et al., 2021). In addition, perhaps also concerning and perplexing is that astrocytes seem not to be activated by fulminant a SARS-CoV-2 brain infection (at least in mice), with their antiviral and neuroprotective activities thus presumably largely absent.

## MATERIALS AND METHODS

### Ethics statements and regulatory compliance

All mouse work was conducted in accordance with the Australian code for the care and use of animals for scientific purposes (National Health and Medical Research Council, Australia). Mouse work was approved by the QIMR Berghofer MRI Animal Ethics Committee (P3600). All infectious SARS-CoV-2 work was conducted in the BioSafety Level 3 (PC3) facility at the QIMR Berghofer MRI (Department of Agriculture, Fisheries and Forestry, certification Q2326 and Office of the Gene Technology Regulator certification 3445). Breeding and use of GM mice was approved under a Notifiable Low Risk Dealing (NLRD) Identifier: NLRD_Suhrbier_Oct2020: NLRD 1.1(a). Mice were euthanized using carbon dioxide.

Collection of nasal swabs from consented COVID-19 patients was approved by the QIMR Berghofer Medical Research Institute Human Research Ethics Committee (P3600) and by the University of Queensland HREC (2022/HE001492).

### SARS-CoV-2 isolates

An original (ancestral) ancestral isolate, SARS-CoV-2_QLD02_ (hCoV-19/Australia/QLD02/2020) (GISAID accession EPI_ISL_407896) was kindly provided by Dr Alyssa Pyke and Fredrick Moore (Queensland Health Forensic & Scientific Services, Queensland Department of Health, Brisbane, Australia) (Rawle et al., 2021). BA.1 and BA.5 omicron isolates were obtained at QIMR Berghofer MRI from nasal swabs from consented COVID-19 patients (Morgan et al., 2023; Yan et al., 2022) that were seeded onto Vero E6 cells (ATCC C1008). Infected Vero E6 cells analysed by RNA-Seq and viral genomes *de novo* assembled using Trinity v 2.8.4. The omicron BA.1 isolate, SARS-CoV-2_QIMR01_ (SARS-CoV-2/human/AUS/QIMR03/2022), belongs to the BA.1.17 lineage (GenBank: ON819429 and GISAID EPI_ISL_13414183) (Morgan et al., 2023; Yan et al., 2022). The omicron BA.5 isolate, SARS-CoV-2_QIMR03_ (SARS-CoV-2/human/AUS/QIMR03/2022) belongs to the BE.1 sublineage (GenBank: OP604184.1). The XBB isolate (SARS-CoV-2_UQ01_) was obtained from nasopharyngeal aspirates of consented COVID-19 patient at the University of Queensland using Vero E6-TMPRSS2 cells (Amarilla et al., 2021). The isolate (deposited as hCoV-19/Australia/UQ01/2023; GISAID EPI_ISL_17784860) is XBB.1.9.2.1.4 (Pango EG.1.4) a recombinant of BA.2.10.1 and BA.2.75. Virus stocks were propagated in Vero E6 cells, viral stocks and tissue culture supernatants were checked for endotoxin (Johnson et al., 2005) and mycoplasma (MycoAlert, Lonza) (La Linn et al., 1995). Viruses were titered using CCID_50_ assays (Yan et al., 2021).

### Mouse infection and monitoring

K18-hACE2 mice (strain B6.Cg-Tg(K18-ACE2)2Prlmn/J, JAX Stock No: 034860) were purchased from The Jackson Laboratory, USA, and were maintained in-house as heterozygotes by backcrossing to C57BL/6J mice (Animal Resources Center, Canning Vale WA, Australia) as described (Bishop et al., 2022). Heterozygotes were inter-crossed to generate a homozygous K18-hACE2 transgenic mouse line. Genotyping was undertaken by PCR and sequencing across a SNP that associates with the hACE2 transgene to distinguish heterozygotes (TTTG(A/C)AAAC) from homozygotes (TTTGCAAAC). The mice were held under standard animal house conditions (for details see (Yan et al., 2022)) and female mice (≈10-20 weeks of age) received intrapulmonary infections delivered via the intranasal route with 5×10^4^ CCID_50_ of virus in 50 μl RPMI 1640 while under light anaesthesia as described (Dumenil et al., 2022). Each group of mice within an experiment had a similar age range and distribution, with the mean age for each group not differing by more than 1 week. Mice were weighed and overt disease symptoms scored as described (Dumenil et al., 2022). Mice were euthanized using CO_2_, and tissue titers determined using CCID_50_ assays and Vero E6 cells as described (Dumenil et al., 2022; Rawle et al., 2021).

### Maintenance and expansion of human induced pluripotent stem cells

The human-induced pluripotent cells (hiPSCs) used in this study were reprogrammed from adult dermal fibroblasts (HDFa, Gibco, C0135C) using the CytoTune-iPS 2.0 Sendai Reprogramming Kit (Invitrogen, A16518) (Oikari et al., 2020). They were cultured on human recombinant vitronectin (Thermo Fisher Scientific) coated plates in StemFlex medium (Thermo Fisher Scientific) according to the manufacturer’s guidelines.

### Generation of human cortical organoids

On dayLJ0 of organoid culture, hiPSCs (less than passage 50) were dissociated with StemPro Accutase (Thermo Fisher Scientific) to generate a cell suspension. Cells were plated 5000/well into an ultra-low-binding 96-well plate (Corning) in StemFlex media supplemented with 10 μM ROCK inhibitor Y-27632 (STEMCELL Technologies, Vancouver, Canada). From days 1-5, media was changed daily with StemFlex medium supplemented with 2 μM Dorsomorphine (Abcam) and 10 μM SB-431542 (Stemcell technologies). On day 5, the medium was replaced with a Neuro-induction medium consisting of DMEM/F12 (Thermo Fisher Scientific), 1% N2 Supplement (Thermo Fisher Scientific), 10 μg/ml heparin (STEMCELL Technologies), 1% penicillin/streptomycin (Thermo Fisher Scientific), 1% Non-essential Amino Acids (Thermo Fisher Scientific), 1% glutamax (Thermo Fisher Scientific) and 10 ng/ml FGF2 (Stemcell Technologies). On day 7, organoids were embedded in Matrigel (Corning), transferred to an ultra-low-binding 24-well plate (Corning) (one organoid per well), and continued to grow in Neuroinduction medium for three more days. On day 10, organoids were supplemented with differentiation medium, consisting of Neurobasal medium, 1% N2, 2% B27 supplements (Thermo Fisher Scientific), 0.5% Penicillin/Streptomycin, 1% Glutamax, 1% Non-essential Amino Acids, 50 µM of 2-mercaptoenthanol (Merck), 2.5 μg/ml Insulin (Merck), 1% Knockout Serum Replacement (Thermo Fisher Scientific), 10 ng/ml FGF2, 1 µM CHIR99021 (Stemcell Technologies) and placed in a CelVivo Clinostar incubator (Invitro Technologies) (24 organoids per Clinoreactor) spinning at 20 rpm. All media changes from 10 days onwards were performed every other day.

### Infection of cortical brain organoids

Organoids (30 days old) were transferred from each Clinoreactor into an ultra-low-binding 24-well plate (one organoid per well), infected with various SARS-CoV-2 strains at a multiplicity of infection (MOI) of 1 and placed within a humidified tissue culture incubator at 37C, 5% CO_2_ for 2 hours. Organoids were then washed twice with media, transferred into 50 mm LUMOX gas exchange dishes (SARSTEDT) (4 organoids per dish) containing 7 ml of differentiation media, and placed within a humidified tissue culture incubator at 37°C, 5% CO_2_ for 4 days.

### Histology and immunohistochemistry

H&E staining of formalin fixed paraffin wax embedded tissue sections was undertaken as described (Amarilla et al., 2021; Rawle et al., 2021). Immunohistochemistry was undertaken as described using the in-house developed anti-spike monoclonal antibody, SCV-1E8 (Morgan et al., 2023).

### Dual labelling fluorescence immunohistochemistry

Paraffin embedded K18-hACE2 mouse brains were sectioned on a rotary microtome into 10 μm sagittal sections. Sections were dewaxed and antigen retrieval was performed in Antigen Recovery Solution (citrate buffer solution consisting of 10 mM sodium citrate, 0.05% SDS 0.01%, pH 6.0) at 90°C for 10 min in a BioCare decloaker. Slides were blocked with blocking buffer (0.5% BSA, 0.05% Saponin, 0.01% Triton X-100, 0.05% Sodium Azide in 0.1M Phosphate Buffered Saline (PBS) for 30 min at room temperature and subsequently incubated in primary antibody diluted in blocking buffer at room temperature for three days in a humidity chamber using anti-S-protein (mouse, 1:100, in-house antibody SCV-1E8 (Morgan et al., 2023)) in conjunction with either anti-NeuN (chicken, 1:3000, Merck, ABN91), anti-Iba1 (rabbit, 1:1000, Wako, #019-19741), or anti-GFAP (rabbit, 1:1000, Abcam, ab7260). Slides were washed 4 times in PBS for 15 min. Slides were further incubated in blocking buffer for 5 min prior to adding the species-specific secondary antibody at room temperature overnight in a light-proof humidity chamber: Alexa fluor-546 anti-mouse (1:1000, Invitrogen, A11030), Alexa fluor-647 anti-rabbit (1:1000, Invitrogen, A32733), or Alexa fluor-488 anti-chicken (1:1000, Invitrogen, A11039). Slides were washed once in PBS and incubated in DAPI diluted (1 μg/ ml, Merck) in saline for 5 min. Slides were washed 3x in PBS for 15 min and were mounted with DABCO mounting media.

Sections were imaged using an Olympus UPLXAPO 10x/0.4 NA air objective, 20x/0.8 NA air objective and an UPLXAPO 60x/1.42 NA oil-immersion objective mounted on a spinning disk confocal microscope (SpinSR10; Olympus, Japan) built on an Olympus IX3 body equipped with two ORCA-Fusion BT sCMOS cameras (Hamamatsu Photonics K.K., Japan) and controlled by Olympus cellSens software. Images were acquired as 3D Z-stack tile images and were deconvolved using Huygens Professional Deconvolution Software (Scientific Volume Imaging, Netherlands).

### RNA-Seq and bioinformatics

RNA-Seq was undertaken as described using Illumina Nextseq 550 platform generating 75 bp paired end reads (Bishop et al., 2022; Rawle et al., 2021). The per base sequence quality for >90% bases was above Q30 for all samples. Mouse RNA-Seq reads were aligned to a combined mouse (GRCm39, version M27) and SARS-CoV-2 (Wuhan, NC_045512.2) reference genome using STAR aligner. Organoid RNA-Seq reads were aligned in the same manner except that the human (GRCh38, version 38) reference genome was used. Read counts for host genes and SARS-CoV-2 genomes were generated using RSEM v1.3.1, and differentially expressed genes were determined using EdgeR v3.36.0. To avoid missing type I IFN genes, which have low read counts (Wilson et al., 2017), a low filter of row sum normalized read count >1 was used.

DEGs in direct and indirect interactions were analyzed using Ingenuity Pathway Analysis (IPA, v84978992) (QIAGEN) using the Canonical pathways, Up-Stream Regulators (USR) and Diseases and Functions features as described (Rawle et al., 2022b). Gene Set Enrichment Analyses (GSEAs) were undertaken using GSEA v4.1.0 with gene sets provided in MSigDB (≈ 50,000 gene sets) and in Blood Transcription Modules and log_2_ fold-change-ranked gene lists generated using DESeq2 v1.34.0 as described (Dumenil et al., 2022; Rawle et al., 2022a). Relative abundances of immune cell types in BA.5-infected mouse brains were estimated from RSEM ‘expected counts’ using SpatialDecon v1.4.3 (Danaher et al., 2022) with the ‘ImmuneAtlas_ImmGen_cellFamily’ profile matrix and Pheatmap v1.0.12 in R v4.1.0.

IPA USR cytokine signatures obtained from BA.5-infected K18-hACE2 mouse brains were compared to gene expression data from two studies on COVID-19 patient brains (Fullard et al., 2021; Yang et al., 2021). For the Yang and Fullard studies there were 20 and 45 gene expression data sets, respectively, for the different tissues and cell-types. DEGs sets (n=20 and 45) were derived by applying a q<0.05 filter. These DEG sets were then concatenated to generate a single DEG list for each of the two studies. When a gene was present in more than one DEG set, the mean of the fold changes was used for the concatenated DEG list. IPA USR analysis was then performed as described above.

### Statistics

Statistical analyses of experimental data were performed using IBM SPSS Statistics for Windows, Version 19.0 (IBM Corp., Armonk, NY, USA). The t-test was used when the difference in variances was <4, skewness was >-2 and kurtosis was <2. For non-parametric data the Kolmogorov-Smirnov test was used.

## SUPPLEMENTAL INFORMATION

The paper is supported by supplementary Figures and Tables.

### ACKNOWLEDGEMENTS

The authors thank the following QIMRB staff; Drs Anthony White and Lotta Oikari for providing the iPSC line, Dr I. Anraku for management of the PC3 facility at QIMR Berghofer MRI, Dr Viviana Lutzky for proof reading, Crystal Chang, Clay Winterford and Sang-Hee Park for histology services, the animal house staff for mouse breeding and agistment, and Dr. Gunter Hartel for assistance with statistics. The authors gratefully acknowledge the Advanced Microscopy Facility and the Histology Facility at The Queensland Brain Institute for their support and assistance in this work, Dr Rumelo Amor, Dr Arnaud Gaudin, Dr Andrew Thompson.

## AUTHOR CONTRIBUTIONS

R.S., S.A.E., K.Y., B.T., W.N., and M.L. performed the experiments. T.D. and C.B. performed computational analyses. T.L., R.K.P.S., F.A.M. and D.J.R. contributed to data analysis and interpretation. A.A.K., R.P. and J.D.J.S. supplied vital reagents and intellectual input. A.S. D.J.R and F.A.M. designed the experiments and supervised the study. A.S. wrote the manuscript with input from all the authors.

## DATA AVAILABILITY

All data is provided in the manuscript and accompanying supplementary files. Raw sequencing data (fastq files) generated for this publication for RNA-Seq have been deposited in the NCBI SRA, BioProject: PRJNA813692 and are publicly available

## References

Abbasi, A.Z., Kiyani, D.A., Hamid, S.M., Saalim, M., Fahim, A., and Jalal, N. (2021). Spiking dependence of SARS-CoV-2 pathogenicity on TMPRSS2. Journal of Medical Virology 93, 4205–4218.

Aggarwal, A., Akerman, A., Milogiannakis, V., Silva, M.R., Walker, G., Stella, A.O., Kindinger, A., Angelovich, T., Waring, E., Amatayakul-Chantler, S., et al. (2022). SARS-CoV-2 Omicron BA.5: Evolving tropism and evasion of potent humoral responses and resistance to clinical immunotherapeutics relative to viral variants of concern. eBioMedicine 84, 104270.

Al-Dalahmah, O., Thakur, K.T., Nordvig, A.S., Prust, M.L., Roth, W., Lignelli, A., Uhlemann, A.C., Miller, E.H., Kunnath-Velayudhan, S., Del Portillo, A., et al. (2020). Neuronophagia and microglial nodules in a SARS-CoV-2 patient with cerebellar hemorrhage. Acta Neuropathol Commun 8, 147.

Altmayer, V., Ziveri, J., Frere, C., Salem, J.E., Weiss, N., Cao, A., Marois, C., Rohaut, B., Demeret, S., Bourdoulous, S., et al. (2022). Endothelial cell biomarkers in critically ill COVID-19 patients with encephalitis. J Neurochem 161, 492–505.

Amarilla, A.A., Sng, J.D.J., Parry, R., Deerain, J.M., Potter, J.R., Setoh, Y.X., Rawle, D.J., Le, T.T., Modhiran, N., Wang, X., et al. (2021). A versatile reverse genetics platform for SARS-CoV-2 and other positive-strand RNA viruses. Nat Commun 12, 3431.

Andrews, M.G., Mukhtar, T., Eze, U.C., Simoneau, C.R., Ross, J., Parikshak, N., Wang, S., Zhou, L., Koontz, M., Velmeshev, D., et al. (2022). Tropism of SARS-CoV-2 for human cortical astrocytes. Proc Natl Acad Sci U S A 119, e2122236119.

Arunachalam, P.S., Wimmers, F., Mok, C.K.P., Perera, R., Scott, M., Hagan, T., Sigal, N., Feng, Y., Bristow, L., Tak-Yin Tsang, O., et al. (2020). Systems biological assessment of immunity to mild versus severe COVID-19 infection in humans. Science 369, 1210–1220.

Aschman, T., Mothes, R., Heppner, F.L., and Radbruch, H. (2022). What SARS-CoV-2 does to our brains. Immunity 55, 1159–1172.

Awogbindin, I.O., Ben-Azu, B., Olusola, B.A., Akinluyi, E.T., Adeniyi, P.A., Di Paolo, T., and Tremblay, M.E. (2021). Microglial Implications in SARS-CoV-2 Infection and COVID-19: Lessons From Viral RNA Neurotropism and Possible Relevance to Parkinson’s Disease. Front Cell Neurosci 15, 670298.

Banerjee, A.K., Blanco, M.R., Bruce, E.A., Honson, D.D., Chen, L.M., Chow, A., Bhat, P., Ollikainen, N., Quinodoz, S.A., Loney, C., et al. (2020). SARS-CoV-2 Disrupts Splicing, Translation, and Protein Trafficking to Suppress Host Defenses. Cell 183, 1325–1339 e1321.

Bauer, L., Laksono, B.M., de Vrij, F.M.S., Kushner, S.A., Harschnitz, O., and van Riel, D. (2022a). The neuroinvasiveness, neurotropism, and neurovirulence of SARS-CoV-2. Trends Neurosci 45, 358–368.

Bauer, L., Rissmann, M., Benavides, F.F.W., Leijten, L., van Run, P., Begeman, L., Veldhuis Kroeze, E.J.B., Lendemeijer, B., Smeenk, H., de Vrij, F.M.S., et al. (2022b). In vitro and in vivo differences in neurovirulence between D614G, Delta And Omicron BA.1 SARS-CoV-2 variants. Acta Neuropathologica Communications 10, 124.

Beckman, D., Bonillas, A., Diniz, G.B., Ott, S., Roh, J.W., Elizaldi, S.R., Schmidt, B.A., Sammak, R.L., Van Rompay, K.K.A., Iyer, S.S., et al. (2022). SARS-CoV-2 infects neurons and induces neuroinflammation in a non-human primate model of COVID-19. Cell Rep 41, 111573.

Bellucci, G., Rinaldi, V., Buscarinu, M.C., Renie, R., Bigi, R., Pellicciari, G., Morena, E., Romano, C., Marrone, A., Mechelli, R., et al. (2021). Multiple Sclerosis and SARS-CoV-2: Has the Interplay Started? Front Immunol 12, 755333.

Bishop, C.R., Dumenil, T., Rawle, D.J., Le, T.T., Yan, K., Tang, B., Hartel, G., and Suhrbier, A. (2022). Mouse models of COVID-19 recapitulate inflammatory pathways rather than gene expression. PLoS Pathog 18, e1010867.

Bowe, B., Xie, Y., and Al-Aly, Z. (2022). Acute and postacute sequelae associated with SARS-CoV-2 reinfection. Nat Med 28, 2398–2405.

Branche, A.R., Rouphael, N.G., Diemert, D.J., Falsey, A.R., Losada, C., Baden, L.R., Frey, S.E., Whitaker, J.A., Little, S.J., Anderson, E.J., et al. (2022). SARS-CoV-2 Variant Vaccine Boosters Trial: Preliminary Analyses. medRxiv.

Bulfamante, G., Chiumello, D., Canevini, M.P., Priori, A., Mazzanti, M., Centanni, S., and Felisati, G. (2020). First ultrastructural autoptic findings of SARS -Cov-2 in olfactory pathways and brainstem. Minerva Anestesiol 86, 678–679.

Bullen, C.K., Hogberg, H.T., Bahadirli-Talbott, A., Bishai, W.R., Hartung, T., Keuthan, C., Looney, M.M., Pekosz, A., Romero, J.C., Sille, F.C.M., et al. (2020). Infectability of human BrainSphere neurons suggests neurotropism of SARS-CoV-2. ALTEX 37, 665–671.

Burks, S.M., Rosas-Hernandez, H., Alejandro Ramirez-Lee, M., Cuevas, E., and Talpos, J.C. (2021). Can SARS-CoV-2 infect the central nervous system via the olfactory bulb or the blood-brain barrier? Brain Behav Immun 95, 7–14.

Cantuti-Castelvetri, L., Ojha, R., Pedro, L.D., Djannatian, M., Franz, J., Kuivanen, S., van der Meer, F., Kallio, K., Kaya, T., Anastasina, M., et al. (2020). Neuropilin-1 facilitates SARS-CoV-2 cell entry and infectivity. Science 370, 856–860.

Cao, Y., Yisimayi, A., Jian, F., Song, W., Xiao, T., Wang, L., Du, S., Wang, J., Li, Q., Chen, X., et al. (2022). BA.2.12.1, BA.4 and BA.5 escape antibodies elicited by Omicron infection. Nature 608, 593–602.

Carabelli, A.M., Peacock, T.P., Thorne, L.G., Harvey, W.T., Hughes, J., Consortium, C.-G.U., Peacock, S.J., Barclay, W.S., de Silva, T.I., Towers, G.J., et al. (2023). SARS-CoV-2 variant biology: immune escape, transmission and fitness. Nat Rev Microbiol 21, 162–177.

Carossino, M., Kenney, D., O’Connell, A.K., Montanaro, P., Tseng, A.E., Gertje, H.P., Grosz, K.A., Ericsson, M., Huber, B.R., Kurnick, S.A., et al. (2022). Fatal Neurodissemination and SARS-CoV-2 Tropism in K18-hACE2 Mice Is Only Partially Dependent on hACE2 Expression. In Viruses, pp. 535.

Chakraborty, S., and Basu, A. (2022). Catching hold of COVID-19-related encephalitis by tracking ANGPTL4 signature in blood: An Editorial Highlight for “Endothelial cell biomarkers in critically ill COVID-19-patients with encephalitis”: An Editorial Highlight for “Endothelial cell biomarkers in critically ill COVID-19-patients with encephalitis” on page 492. J Neurochem 161, 458–462.

Chen, C.S., Chang, C.N., Hu, C.F., Jian, M.J., Chung, H.Y., Chang, C.K., Perng, C.L., Hung, K.S., Chang, F.Y., Wang, C.H., et al. (2022). Critical pediatric neurological illness associated with COVID-19 (Omicron BA.2.3.7 variant) infection in Taiwan: immunological assessment and viral genome analysis in tertiary medical center. Int J Infect Dis 124, 45–48.

Chen, D.Y., Chin, C.V., Kenney, D., Tavares, A.H., Khan, N., Conway, H.L., Liu, G., Choudhary, M.C., Gertje, H.P., O’Connell, A.K., et al. (2023). Spike and nsp6 are key determinants of SARS-CoV-2 Omicron BA.1 attenuation. Nature 615, 143–150.

Cloete, J., Kruger, A., Masha, M., du Plessis, N.M., Mawela, D., Tshukudu, M., Manyane, T., Komane, L., Venter, M., Jassat, W., et al. (2022). Paediatric hospitalisations due to COVID-19 during the first SARS-CoV-2 omicron (B.1.1.529) variant wave in South Africa: a multicentre observational study. Lancet Child Adolesc Health 6, 294–302.

Crunfli, F., Carregari, V.C., Veras, F.P., Silva, L.S., Nogueira, M.H., Antunes, A., Vendramini, P.H., Valenca, A.G.F., Brandao-Teles, C., Zuccoli, G.D.S., et al. (2022). Morphological, cellular, and molecular basis of brain infection in COVID-19 patients. Proc Natl Acad Sci U S A 119, e2200960119.

Danaher, P., Kim, Y., Nelson, B., Griswold, M., Yang, Z., Piazza, E., and Beechem, J.M. (2022). Advances in mixed cell deconvolution enable quantification of cell types in spatial transcriptomic data. Nat Commun 13, 385.

Dang, T.Q., La, D.T., and Tran, T.N. (2022). Myeloencephalitis as the only presentation of Omicron SARS-CoV-2 infection. BMJ Case Rep 15, e251922.

Davies, M.A., Morden, E., Rousseau, P., Arendse, J., Bam, J.L., Boloko, L., Cloete, K., Cohen, C., Chetty, N., Dane, P., et al. (2023). Outcomes of laboratory-confirmed SARS-CoV-2 infection during resurgence driven by Omicron lineages BA.4 and BA.5 compared with previous waves in the Western Cape Province, South Africa. Int J Infect Dis 127, 63–68.

Davis, H.E., McCorkell, L., Vogel, J.M., and Topol, E.J. (2023). Long COVID: major findings, mechanisms and recommendations. Nat Rev Microbiol 21, 133–146.

De Hert, M., Mazereel, V., Stroobants, M., De Picker, L., Van Assche, K., and Detraux, J. (2021). COVID-19-Related Mortality Risk in People With Severe Mental Illness: A Systematic and Critical Review. Front Psychiatry 12, 798554.

de Paula, J.J., Paiva, R., Souza-Silva, N.G., Rosa, D.V., Duran, F.L.S., Coimbra, R.S., Costa, D.S., Dutenhefner, P.R., Oliveira, H.S.D., Camargos, S.T., et al. (2023). Selective visuoconstructional impairment following mild COVID-19 with inflammatory and neuroimaging correlation findings. Mol Psychiatry 28, 553–563.

Dong, W., Mead, H., Tian, L., Park, J.G., Garcia, J.I., Jaramillo, S., Barr, T., Kollath, D.S., Coyne, V.K., Stone, N.E., et al. (2022). The K18-Human ACE2 Transgenic Mouse Model Recapitulates Non-severe and Severe COVID-19 in Response to an Infectious Dose of the SARS-CoV-2 Virus. J Virol 96, e0096421.

Douaud, G., Lee, S., Alfaro-Almagro, F., Arthofer, C., Wang, C., McCarthy, P., Lange, F., Andersson, J.L.R., Griffanti, L., Duff, E., et al. (2022). SARS-CoV-2 is associated with changes in brain structure in UK Biobank. Nature 604, 697–707.

Du, P., Gao, G.F., and Wang, Q. (2022). The mysterious origins of the Omicron variant of SARS-CoV-2. Innovation (Camb) 3, 100206.

Dumenil, T., Le, T.T., Rawle, D.J., Yan, K., Tang, B., Nguyen, W., Bishop, C., and Suhrbier, A. (2022). Warmer ambient air temperatures reduce nasal turbinate and brain infection, but increase lung inflammation in the K18-hACE2 mouse model of COVID-19. Sci Total Environ 859, 160163.

Durante, M.A., Kurtenbach, S., Sargi, Z.B., Harbour, J.W., Choi, R., Kurtenbach, S., Goss, G.M., Matsunami, H., and Goldstein, B.J. (2020). Single-cell analysis of olfactory neurogenesis and differentiation in adult humans. Nat Neurosci 23, 323–326.

El-Kassas, M., Alboraie, M., Elbadry, M., El Sheemy, R., Abdellah, M., Afify, S., Madkour, A., Zaghloul, M., Awad, A., Wifi, M.N., et al. (2023). Non-pulmonary involvement in COVID-19: A systemic disease rather than a pure respiratory infection. World J Clin Cases 11, 493–505.

Emmi, A., Rizzo, S., Barzon, L., Sandre, M., Carturan, E., Sinigaglia, A., Riccetti, S., Della Barbera, M., Boscolo-Berto, R., Cocco, P., et al. (2023). Detection of SARS-CoV-2 viral proteins and genomic sequences in human brainstem nuclei. npj Parkinson’s Disease 9, 25.

Fernandez-Castaneda, A., Lu, P., Geraghty, A.C., Song, E., Lee, M.H., Wood, J., O’Dea, M.R., Dutton, S., Shamardani, K., Nwangwu, K., et al. (2022). Mild respiratory COVID can cause multi-lineage neural cell and myelin dysregulation. Cell 185, 2452–2468 e2416.

Ferren, M., Favede, V., Decimo, D., Iampietro, M., Lieberman, N.A.P., Weickert, J.L., Pelissier, R., Mazelier, M., Terrier, O., Moscona, A., et al. (2021). Hamster organotypic modeling of SARS-CoV-2 lung and brainstem infection. Nat Commun 12, 5809.

Ferrucci, R., Cuffaro, L., Capozza, A., Rosci, C., Maiorana, N., Groppo, E., Reitano, M.R., Poletti, B., Ticozzi, N., Tagliabue, L., et al. (2023). Brain positron emission tomography (PET) and cognitive abnormalities one year after COVID-19. J Neurol.

Fodoulian, L., Tuberosa, J., Rossier, D., Boillat, M., Kan, C., Pauli, V., Egervari, K., Lobrinus, J.A., Landis, B.N., Carleton, A., et al. (2020). SARS-CoV-2 Receptors and Entry Genes Are Expressed in the Human Olfactory Neuroepithelium and Brain. iScience 23, 101839.

Fullard, J.F., Lee, H.C., Voloudakis, G., Suo, S., Javidfar, B., Shao, Z., Peter, C., Zhang, W., Jiang, S., Corvelo, A., et al. (2021). Single-nucleus transcriptome analysis of human brain immune response in patients with severe COVID-19. Genome Med 13, 118.

Fumagalli, V., Rava, M., Marotta, D., Di Lucia, P., Laura, C., Sala, E., Grillo, M., Bono, E., Giustini, L., Perucchini, C., et al. (2022). Administration of aerosolized SARS-CoV-2 to K18-hACE2 mice uncouples respiratory infection from fatal neuroinvasion. Sci Immunol 7, eabl9929.

Graham, E.L., Koralnik, I.J., and Liotta, E.M. (2022). Therapeutic Approaches to the Neurologic Manifestations of COVID-19. Neurotherapeutics 19, 1435–1466.

Guimond, S.E., Mycroft-West, C.J., Gandhi, N.S., Tree, J.A., Le, T.T., Spalluto, C.M., Humbert, M.V., Buttigieg, K.R., Coombes, N., Elmore, M.J., et al. (2022). Synthetic Heparan Sulfate Mimetic Pixatimod (PG545) Potently Inhibits SARS-CoV-2 by Disrupting the Spike-ACE2 Interaction. ACS Cent Sci 8, 527–545.

Guo, K., Barrett, B.S., Morrison, J.H., Mickens, K.L., Vladar, E.K., Hasenkrug, K.J., Poeschla, E.M., and Santiago, M.L. (2022). Interferon resistance of emerging SARS-CoV-2 variants. Proc Natl Acad Sci U S A 119, e2203760119.

Halfmann, P.J., Iida, S., Iwatsuki-Horimoto, K., Maemura, T., Kiso, M., Scheaffer, S.M., Darling, T.L., Joshi, A., Loeber, S., Singh, G., et al. (2022). SARS-CoV-2 Omicron virus causes attenuated disease in mice and hamsters. Nature 603, 687–692.

Hansen, C.H., Friis, N.U., Bager, P., Stegger, M., Fonager, J., Fomsgaard, A., Gram, M.A., Christiansen, L.E., Ethelberg, S., Legarth, R., et al. (2023). Risk of reinfection, vaccine protection, and severity of infection with the BA.5 omicron subvariant: a nation-wide population-based study in Denmark. The Lancet Infectious Diseases 23, 167–176.

Hosseinzadeh, H., and Goldschneider, I. (1993). Recent thymic emigrants in the rat express a unique antigenic phenotype and undergo post-thymic maturation in peripheral lymphoid tissues. J Immunol 150, 1670–1679.

Hou, Y., Li, C., Yoon, C., Leung, O.W., You, S., Cui, X., Chan, J.F.-W., Pei, D., Cheung, H.H., and Chu, H. (2022). Enhanced replication of SARS-CoV-2 Omicron BA.2 in human forebrain and midbrain organoids. Signal Transduction and Targeted Therapy 7, 381.

Houben, E., Hellings, N., and Broux, B. (2019). Oncostatin M, an Underestimated Player in the Central Nervous System. Front Immunol 10, 1165.

Islam, M.A., Cavestro, C., Alam, S.S., Kundu, S., Kamal, M.A., and Reza, F. (2022). Encephalitis in Patients with COVID-19: A Systematic Evidence-Based Analysis. Cells 11, 2575.

Islam, M.A., Kaifa, F.H., Chandran, D., Bhattacharya, M., Chakraborty, C., Bhattacharya, P., and Dhama, K. (2023). XBB.1.5: A new threatening SARS-CoV-2 Omicron subvariant. Front Microbiol 14, 1154296.

Ismail, II, and Salama, S. (2022). Association of CNS demyelination and COVID-19 infection: an updated systematic review. J Neurol 269, 541–576.

Iwata-Yoshikawa, N., Kakizaki, M., Shiwa-Sudo, N., Okura, T., Tahara, M., Fukushi, S., Maeda, K., Kawase, M., Asanuma, H., Tomita, Y., et al. (2022). Essential role of TMPRSS2 in SARS-CoV-2 infection in murine airways. Nature Communications 13, 6100.

Jiao, L., Yang, Y., Yu, W., Zhao, Y., Long, H., Gao, J., Ding, K., Ma, C., Li, J., Zhao, S., et al. (2021). The olfactory route is a potential way for SARS-CoV-2 to invade the central nervous system of rhesus monkeys. Signal Transduct Target Ther 6, 169.

Johnson, B.J., Le, T.T., Dobbin, C.A., Banovic, T., Howard, C.B., Flores Fde, M., Vanags, D., Naylor, D.J., Hill, G.R., and Suhrbier, A. (2005). Heat shock protein 10 inhibits lipopolysaccharide-induced inflammatory mediator production. J Biol Chem 280, 4037–4047.

Kang, S.W., Park, H., Kim, J.Y., Lim, S.Y., Lee, S., Bae, J.Y., Kim, J., Chang, E., Bae, S., Jung, J., et al. (2023). Comparison of the clinical and virological characteristics of SARS-CoV-2 Omicron BA.1/BA.2 and omicron BA.5 variants: A prospective cohort study. J Infect.

Karpiel, I., Starcevic, A., and Urzeniczok, M. (2022). Database and AI Diagnostic Tools Improve Understanding of Lung Damage, Correlation of Pulmonary Disease and Brain Damage in COVID-19. Sensors (Basel) 22, 6312.

Kettunen, P., Lesnikova, A., Rasanen, N., Ojha, R., Palmunen, L., Laakso, M., Lehtonen, S., Kuusisto, J., Pietilainen, O., Saber, S.H., et al. (2023). SARS-CoV-2 Infection of Human Neurons Is TMPRSS2 Independent, Requires Endosomal Cell Entry, and Can Be Blocked by Inhibitors of Host Phosphoinositol-5 Kinase. J Virol 97, e0014423.

Kimura, I., Yamasoba, D., Tamura, T., Nao, N., Suzuki, T., Oda, Y., Mitoma, S., Ito, J., Nasser, H., Zahradnik, J., et al. (2022). Virological characteristics of the SARS-CoV-2 Omicron BA.2 subvariants, including BA.4 and BA.5. Cell 185, 3992–4007 e3916.

Kong, W., Montano, M., Corley, M.J., Helmy, E., Kobayashi, H., Kinisu, M., Suryawanshi, R., Luo, X., Royer, L.A., Roan, N.R., et al. (2022). Neuropilin-1 Mediates SARS-CoV-2 Infection of Astrocytes in Brain Organoids, Inducing Inflammation Leading to Dysfunction and Death of Neurons. mBio 13, e0230822.

Kouamen, A.C., Da Cruz, H., Hamidouche, M., Lamy, A., Lloyd, A., Castro Alvarez, J., Roussel, M., Josset, L., Enouf, V., Felici, C., et al. (2022). Rapid investigation of BA.4/BA.5 cases in France. Front Public Health 10, 1006631.

Kumari, P., Rothan, H.A., Natekar, J.P., Stone, S., Pathak, H., Strate, P.G., Arora, K., Brinton, M.A., and Kumar, M. (2021). Neuroinvasion and Encephalitis Following Intranasal Inoculation of SARS-CoV-2 in K18-hACE2 Mice. Viruses 13, 132.

La Linn, M., Bellett, A.J., Parsons, P.G., and Suhrbier, A. (1995). Complete removal of mycoplasma from viral preparations using solvent extraction. J Virol Methods 52, 51–54.

Ledford, H. (2022). Severe COVID could cause markers of old age in the brain. Nature 612, 389.

Li, K., Meyerholz David, K., Bartlett Jennifer, A., and McCray Paul, B. (2021). The TMPRSS2 Inhibitor Nafamostat Reduces SARS-CoV-2 Pulmonary Infection in Mouse Models of COVID-19. mBio 12, e00970–00921.

Ludvigsson, J.F. (2022). Convulsions in children with COVID-19 during the Omicron wave. Acta Paediatr 111, 1023–1026.

Maiese, A., Manetti, A.C., Bosetti, C., Del Duca, F., La Russa, R., Frati, P., Di Paolo, M., Turillazzi, E., and Fineschi, V. (2021). SARS-CoV-2 and the brain: A review of the current knowledge on neuropathology in COVID-19. Brain Pathol 31, e13013.

Mallapaty, S. (2022). Where did Omicron come from? Three key theories. Nature 602, 26–28.

Martin, M., Paes, V.R., Cardoso, E.F., Neto, C., Kanamura, C.T., Leite, C.D.C., Otaduy, M.C.G., Monteiro, R.A.A., Mauad, T., da Silva, L.F.F., et al. (2022). Postmortem brain 7T MRI with minimally invasive pathological correlation in deceased COVID-19 subjects. Insights Imaging 13, 7.

Martínez-Mármol, R., Giordano-Santini, R., Kaulich, E., Cho, A.-N., Przybyla, M., Riyadh, M.A., Robinson, E., Chew, K.Y., Amor, R., Meunier, F.A., et al. (2023). SARS-CoV-2 infection and viral fusogens cause neuronal and glial fusion that compromises neuronal activity. Science Advances 9, eadg2248.

Matschke, J., Lutgehetmann, M., Hagel, C., Sperhake, J.P., Schroder, A.S., Edler, C., Mushumba, H., Fitzek, A., Allweiss, L., Dandri, M., et al. (2020). Neuropathology of patients with COVID-19 in Germany: a post-mortem case series. Lancet Neurol 19, 919–929.

Meinhardt, J., Radke, J., Dittmayer, C., Franz, J., Thomas, C., Mothes, R., Laue, M., Schneider, J., Brunink, S., Greuel, S., et al. (2021). Olfactory transmucosal SARS-CoV-2 invasion as a port of central nervous system entry in individuals with COVID-19. Nat Neurosci 24, 168–175.

Meng, B., Abdullahi, A., Ferreira, I.A.T.M., Goonawardane, N., Saito, A., Kimura, I., Yamasoba, D., Gerber, P.P., Fatihi, S., Rathore, S., et al. (2022). Altered TMPRSS2 usage by SARS-CoV-2 Omicron impacts infectivity and fusogenicity. Nature 603, 706–714.

Mesci, P., de Souza, J.S., Martin-Sancho, L., Macia, A., Saleh, A., Yin, X., Snethlage, C., Adams, J.W., Avansini, S.H., Herai, R.H., et al. (2022). SARS-CoV-2 infects human brain organoids causing cell death and loss of synapses that can be rescued by treatment with Sofosbuvir. PLOS Biology 20, e3001845.

Monje, M., and Iwasaki, A. (2022). The neurobiology of long COVID. Neuron 110, 3484–3496.

Morgan, M.S., Yan, K., Le, T.T., Johnston, R.A., Amarilla, A.A., Muller, D.A., McMillan, C.L.D., Modhiran, N., Watterson, D., Potter, J.R., et al. (2023). Monoclonal Antibodies Specific for SARS-CoV-2 Spike Protein Suitable for Multiple Applications for Current Variants of Concern. Viruses 15, 139.

Msemburi, W., Karlinsky, A., Knutson, V., Aleshin-Guendel, S., Chatterji, S., and Wakefield, J. (2023). The WHO estimates of excess mortality associated with the COVID-19 pandemic. Nature 613, 130–137.

Mukerji, S.S., and Solomon, I.H. (2021). What can we learn from brain autopsies in COVID-19? Neurosci Lett 742, 135528.

Nchioua, R., Diofano, F., Noettger, S., von Maltitz, P., Stenger, S., Zech, F., Munch, J., Sparrer, K.M.J., Just, S., and Kirchhoff, F. (2022). Strong attenuation of SARS-CoV-2 Omicron BA.1 and increased replication of the BA.5 subvariant in human cardiomyocytes. Signal Transduct Target Ther 7, 395.

Normandin, E., Valizadeh, N., Rudmann, E.A., Uddin, R., Dobbins, S.T., MacInnis, B.L., Padera, R.F., Siddle, K.J., Lemieux, J.E., Sabeti, P.C., et al. (2023). Neuropathological features of SARS-CoV-2 delta and omicron variants. J Neuropathol Exp Neurol, nlad015.

Oikari, L.E., Pandit, R., Stewart, R., Cuní-López, C., Quek, H., Sutharsan, R., Rantanen, L.M., Oksanen, M., Lehtonen, S., de Boer, C.M., et al. (2020). Altered Brain Endothelial Cell Phenotype from a Familial Alzheimer Mutation and Its Potential Implications for Amyloid Clearance and Drug Delivery. Stem Cell Reports 14, 924–939.

Okrzeja, J., Garkowski, A., Kubas, B., and Moniuszko-Malinowska, A. (2023). Imaging and neuropathological findings in patients with Post COVID-19 Neurological Syndrome-A review. Front Neurol 14, 1136348.

Olivarria Gema, M., Cheng, Y., Furman, S., Pachow, C., Hohsfield Lindsay, A., Smith-Geater, C., Miramontes, R., Wu, J., Burns Mara, S., Tsourmas Kate, I., et al. (2022). Microglia Do Not Restrict SARS-CoV-2 Replication following Infection of the Central Nervous System of K18-Human ACE2 Transgenic Mice. Journal of Virology 96, e01969–01921.

Oner, E., Al-Khafaji, K., Mezher, M.H., Demirhan, I., Suhail Wadi, J., Belge Kurutas, E., Yalin, S., and Choowongkomon, K. (2022). Investigation of berberine and its derivatives in Sars Cov-2 main protease structure by molecular docking, PROTOX-II and ADMET methods: in machine learning and in silico study. J Biomol Struct Dyn, 1–16.

Ong, C.P., Ye, Z.-W., Tang, K., Liang, R., Xie, Y., Zhang, H., Qin, Z., Sun, H., Wang, T.-Y., Cheng, Y., et al. (2023). Comparative analysis of SARS-CoV-2 Omicron BA.2.12.1 and BA.5.2 variants. Journal of Medical Virology 95, e28326.

OurWorldinData (2022). SARS-CoV-2 sequences by variant, Jan 30, 2023. https://ourworldindata.org/grapher/covid-variants-bar?country=CAN∼BWA∼ESP∼ZAF∼AUS∼GBR∼USA∼DEU∼ITA∼BEL∼FRA. Accessed June 2023.

Pajo, A.T., Espiritu, A.I., Apor, A., and Jamora, R.D.G. (2021). Neuropathologic findings of patients with COVID-19: a systematic review. Neurol Sci 42, 1255–1266.

Pelizzari, L., Cazzoli, M., Lipari, S., Lagana, M.M., Cabinio, M., Isernia, S., Pirastru, A., Clerici, M., and Baglio, F. (2022). Mid-term MRI evaluation reveals microstructural white matter alterations in COVID-19 fully recovered subjects with anosmia presentation. Ther Adv Neurol Disord 15, 1–10.

Pinto, M.V., and Fernandes, A. (2020). Microglial Phagocytosis-Rational but Challenging Therapeutic Target in Multiple Sclerosis. Int J Mol Sci 21, 5960.

Qasmieh, S.A., Robertson, M.M., Teasdale, C.A., Kulkarni, S.G., Jones, H.E., McNairy, M., Borrell, L.N., and Nash, D. (2023). The prevalence of SARS-CoV-2 infection and long COVID in U.S. adults during the BA.4/BA.5 surge, June-July 2022. Prev Med 169, 107461.

Qu, P., Evans, J.P., Kurhade, C., Zeng, C., Zheng, Y.M., Xu, K., Shi, P.Y., Xie, X., and Liu, S.L. (2023). Determinants and Mechanisms of the Low Fusogenicity and High Dependence on Endosomal Entry of Omicron Subvariants. mBio 14, e0317622.

RADC (2022). Persistent Loss of Smell Due to COVID-19 Closely Connected to Long-Lasting Cognitive Problems. In Alzheimer’s Association International Conference (San Diego: https://aaic.alz.org/downloads2022/COVID-and-Cognition-News-Release-AAIC2022.pdf).

Radhakrishnan, R.K., and Kandasamy, M. (2022). SARS-CoV-2-Mediated Neuropathogenesis, Deterioration of Hippocampal Neurogenesis and Dementia. Am J Alzheimers Dis Other Demen 37, 1–10.

Rawle, D.J., Dumenil, T., Tang, B., Bishop, C.R., Yan, K., Le, T.T., and Suhrbier, A. (2022a). Microplastic consumption induces inflammatory signatures in the colon and prolongs a viral arthritis. Sci Total Environ 809, 152212.

Rawle, D.J., Le, T.T., Dumenil, T., Bishop, C., Yan, K., Nakayama, E., Bird, P.I., and Suhrbier, A. (2022b). Widespread discrepancy in Nnt genotypes and genetic backgrounds complicates granzyme A and other knockout mouse studies. Elife 11, e70207.

Rawle, D.J., Le, T.T., Dumenil, T., Yan, K., Tang, B., Nguyen, W., Watterson, D., Modhiran, N., Hobson-Peters, J., Bishop, C., et al. (2021). ACE2-lentiviral transduction enables mouse SARS-CoV-2 infection and mapping of receptor interactions. PLoS Pathog 17, e1009723.

Rizvi, Z.A., Adhikari, N., Sharma, K., Sadhu sadhu, S., Dandotiya, J., Khatri, R., Singh, V., Vinayakdas, K.V., Samal, S., Mani, S., et al. (2022). Omicron sub-lineage BA.5 infection causes attenuated pathology and results in robust protection in Omicron recovered hACE2 transgenic mice. Available at SSRN: https://ssrn.com/abstract=4243698 or 10.2139/ssrn.4243698.

Rosu, G.C., Mateescu, V.O., Simionescu, A., Istrate-Ofiteru, A.M., Curca, G.C., Pirici, I., Mogoanta, L., Mindrila, I., Kumar-Singh, S., Hostiuc, S., et al. (2022). Subtle vascular and astrocytic changes in the brain of coronavirus disease 2019 (COVID-19) patients. Eur J Neurol 29, 3676–3692.

Rothan, H.A., Kumari, P., Stone, S., Natekar, J.P., Arora, K., Auroni, T.T., and Kumar, M. (2022). SARS-CoV-2 Infects Primary Neurons from Human ACE2 Expressing Mice and Upregulates Genes Involved in the Inflammatory and Necroptotic Pathways. Pathogens 11, 257.

Rothstein, T.L. (2023). Cortical Grey matter volume depletion links to neurological sequelae in post COVID-19 “long haulers”. BMC Neurol 23, 22.

Russell, S.L., Klaver, B.R.A., Harrigan, S.P., Kamelian, K., Tyson, J., Hoang, L., Taylor, M., Sander, B., Mishra, S., Prystajecky, N., et al. (2023). Clinical severity of Omicron subvariants BA.1, BA.2, and BA.5 in a population-based cohort study in British Columbia, Canada. J Med Virol 95, e28423.

Rutkai, I., Mayer, M.G., Hellmers, L.M., Ning, B., Huang, Z., Monjure, C.J., Coyne, C., Silvestri, R., Golden, N., Hensley, K., et al. (2022). Neuropathology and virus in brain of SARS-CoV-2 infected non-human primates. Nat Commun 13, 1745.

Samudyata, Oliveira, A.O., Malwade, S., Rufino de Sousa, N., Goparaju, S.K., Gracias, J., Orhan, F., Steponaviciute, L., Schalling, M., Sheridan, S.D., et al. (2022). SARS-CoV-2 promotes microglial synapse elimination in human brain organoids. Mol Psychiatry 27, 3939–3950.

Sanabria-Diaz, G., Etter, M.M., Melie-Garcia, L., Lieb, J.M., Psychogios, M.N., Hutter, G., and Granziera, C. (2022). Brain cortical alterations in COVID-19 patients with neurological symptoms. Front Neurosci 16, 992165.

Schmidt, N., Lareau, C.A., Keshishian, H., Ganskih, S., Schneider, C., Hennig, T., Melanson, R., Werner, S., Wei, Y., Zimmer, M., et al. (2021). The SARS-CoV-2 RNA–protein interactome in infected human cells. Nature Microbiology 6, 339–353.

Schwabenland, M., Salie, H., Tanevski, J., Killmer, S., Lago, M.S., Schlaak, A.E., Mayer, L., Matschke, J., Puschel, K., Fitzek, A., et al. (2021). Deep spatial profiling of human COVID-19 brains reveals neuroinflammation with distinct microanatomical microglia-T-cell interactions. Immunity 54, 1594–1610 e1511.

Seehusen, F., Clark, J.J., Sharma, P., Bentley, E.G., Kirby, A., Subramaniam, K., Wunderlin-Giuliani, S., Hughes, G.L., Patterson, E.I., Michael, B.D., et al. (2022). Neuroinvasion and Neurotropism by SARS-CoV-2 Variants in the K18-hACE2 Mouse. Viruses 14, 1020.

Sepehrinezhad, A., Gorji, A., and Sahab Negah, S. (2021). SARS-CoV-2 may trigger inflammasome and pyroptosis in the central nervous system: a mechanistic view of neurotropism. Inflammopharmacology 29, 1049–1059.

Serafini, R.A., Frere, J.J., Zimering, J., Giosan, I.M., Pryce, K.D., Golynker, I., Panis, M., Ruiz, A., tenOever, B., and Zachariou, V. (2022). SARS-CoV-2 Airway Infection Results in Time-dependent Sensory Abnormalities in a Hamster Model. bioRxiv.

Serrano, G.E., Walker, J.E., Tremblay, C., Piras, I.S., Huentelman, M.J., Belden, C.M., Goldfarb, D., Shprecher, D., Atri, A., Adler, C.H., et al. (2022). SARS-CoV-2 Brain Regional Detection, Histopathology, Gene Expression, and Immunomodulatory Changes in Decedents with COVID-19. J Neuropathol Exp Neurol 81, 666–695.

Shen, W.B., Logue, J., Yang, P., Baracco, L., Elahi, M., Reece, E.A., Wang, B., Li, L., Blanchard, T.G., Han, Z., et al. (2022). SARS-CoV-2 invades cognitive centers of the brain and induces Alzheimer’s-like neuropathology. bioRxiv.

Shrestha, L.B., Foster, C., Rawlinson, W., Tedla, N., and Bull, R.A. (2022). Evolution of the SARS-CoV-2 omicron variants BA.1 to BA.5: Implications for immune escape and transmission. Rev Med Virol 32, e2381.

Shuai, H., Chan, J.F., Hu, B., Chai, Y., Yuen, T.T., Yin, F., Huang, X., Yoon, C., Hu, J.C., Liu, H., et al. (2022). Attenuated replication and pathogenicity of SARS-CoV-2 B.1.1.529 Omicron. Nature 603, 693–699.

Sigal, A. (2022). Milder disease with Omicron: is it the virus or the pre-existing immunity? Nat Rev Immunol 22, 69–71.

Silva, R.C., da Rosa, M.M., Leao, H.I., Silva, E.D.L., Ferreira, N.T., Albuquerque, A.P.B., Duarte, G.S., Siqueira, A.M., Pereira, M.C., Rego, M., et al. (2023). Brain damage serum biomarkers induced by COVID-19 in patients from northeast Brazil. J Neurovirol, 1–7.

Siow, I., Lee, K.S., Zhang, J.J.Y., Saffari, S.E., and Ng, A. (2021). Encephalitis as a neurological complication of COVID-19: A systematic review and meta-analysis of incidence, outcomes, and predictors. Eur J Neurol 28, 3491–3502.

Song, E., Zhang, C., Israelow, B., Lu-Culligan, A., Prado, A.V., Skriabine, S., Lu, P., Weizman, O.E., Liu, F., Dai, Y., et al. (2021). Neuroinvasion of SARS-CoV-2 in human and mouse brain. J Exp Med 218, e20202135.

Stein, S.R., Ramelli, S.C., Grazioli, A., Chung, J.-Y., Singh, M., Yinda, C.K., Winkler, C.W., Sun, J., Dickey, J.M., Ylaya, K., et al. (2022). SARS-CoV-2 infection and persistence in the human body and brain at autopsy. Nature 612, 758–763.

Surie, D., Bonnell, L., Adams, K., Gaglani, M., Ginde, A.A., Douin, D.J., Talbot, H.K., Casey, J.D., Mohr, N.M., Zepeski, A., et al. (2022). Effectiveness of Monovalent mRNA Vaccines Against COVID-19-Associated Hospitalization Among Immunocompetent Adults During BA.1/BA.2 and BA.4/BA.5 Predominant Periods of SARS-CoV-2 Omicron Variant in the United States - IVY Network, 18 States, December 26, 2021-August 31, 2022. MMWR Morb Mortal Wkly Rep 71, 1327–1334.

Suryawanshi, R.K., Chen, I.P., Ma, T., Syed, A.M., Brazer, N., Saldhi, P., Simoneau, C.R., Ciling, A., Khalid, M.M., Sreekumar, B., et al. (2022). Limited cross-variant immunity from SARS-CoV-2 Omicron without vaccination. Nature 607, 351–355.

Takashita, E., Yamayoshi, S., Simon, V., van Bakel, H., Sordillo, E.M., Pekosz, A., Fukushi, S., Suzuki, T., Maeda, K., Halfmann, P., et al. (2022). Efficacy of Antibodies and Antiviral Drugs against Omicron BA.2.12.1, BA.4, and BA.5 Subvariants. N Engl J Med 387, 468–470.

Tamura, T., Ito, J., Uriu, K., Zahradnik, J., Kida, I., Anraku, Y., Nasser, H., Shofa, M., Oda, Y., Lytras, S., et al. (2023). Virological characteristics of the SARS-CoV-2 XBB variant derived from recombination of two Omicron subvariants. Nat Commun 14, 2800.

Tang, H., Shao, Y., Huang, Y., Qiao, S., An, J., Yan, R., Zhao, X., Meng, F., Du, X., and Qin, F.X. (2023). Evolutionary characteristics of SARS-CoV-2 Omicron subvariants adapted to the host. Signal Transduct Target Ther 8, 211.

Tanne, J.H. (2022). Covid-19: BA.5 variant is now dominant in US as infections rise. BMJ 378, o1770.

Taquet, M., Sillett, R., Zhu, L., Mendel, J., Camplisson, I., Dercon, Q., and Harrison, P.J. (2022). Neurological and psychiatric risk trajectories after SARS-CoV-2 infection: an analysis of 2-year retrospective cohort studies including 1 284 437 patients. Lancet Psychiatry 9, 815–827.

Tarres-Freixas, F., Trinite, B., Pons-Grifols, A., Romero-Durana, M., Riveira-Munoz, E., Avila-Nieto, C., Perez, M., Garcia-Vidal, E., Perez-Zsolt, D., Munoz-Basagoiti, J., et al. (2022). Heterogeneous Infectivity and Pathogenesis of SARS-CoV-2 Variants Beta, Delta and Omicron in Transgenic K18-hACE2 and Wildtype Mice. Front Microbiol 13, 840757.

The-Jackson-Laboratory. B6.Cg-Tg(K18-ACE2)2Prlmn/J. Protocol 38275. https://www.jax.org/Protocol?stockNumber=034860&protocolID=38275. Accessed June 2023.

Tian, M., Liu, W., Li, X., Zhao, P., Shereen, M.A., Zhu, C., Huang, S., Liu, S., Yu, X., Yue, M., et al. (2021). HIF-1alpha promotes SARS-CoV-2 infection and aggravates inflammatory responses to COVID-19. Signal Transduct Target Ther 6, 308.

Tuekprakhon, A., Huo, J., Nutalai, R., Dijokaite-Guraliuc, A., Zhou, D., Ginn, H.M., Selvaraj, M., Liu, C., Mentzer, A.J., Supasa, P., et al. (2022). Further antibody escape by Omicron BA.4 and BA.5 from vaccine and BA.1 serum. bioRxiv, 2022.2005.2021.492554.

Uraki, R., Halfmann, P.J., Iida, S., Yamayoshi, S., Furusawa, Y., Kiso, M., Ito, M., Iwatsuki-Horimoto, K., Mine, S., Kuroda, M., et al. (2022). Characterization of SARS-CoV-2 Omicron BA.4 and BA.5 isolates in rodents. Nature 612, 540–545.

Vidal, E., Lopez-Figueroa, C., Rodon, J., Perez, M., Brustolin, M., Cantero, G., Guallar, V., Izquierdo-Useros, N., Carrillo, J., Blanco, J., et al. (2022). Chronological brain lesions after SARS-CoV-2 infection in hACE2-transgenic mice. Vet Pathol 59, 613–626.

Wang, Q., Guo, Y., Iketani, S., Nair, M.S., Li, Z., Mohri, H., Wang, M., Yu, J., Bowen, A.D., Chang, J.Y., et al. (2022). Antibody evasion by SARS-CoV-2 Omicron subvariants BA.2.12.1, BA.4 and BA.5. Nature 608, 603–608.

Wilson, J.A.C., Prow, N.A., Schroder, W.A., Ellis, J.J., Cumming, H.E., Gearing, L.J., Poo, Y.S., Taylor, A., Hertzog, P.J., Di Giallonardo, F., et al. (2017). RNA-Seq analysis of chikungunya virus infection and identification of granzyme A as a major promoter of arthritic inflammation. PLOS Pathogens 13, e1006155.

Wolter, N., Jassat, W., Walaza, S., Welch, R., Moultrie, H., Groome, M.J., Amoako, D.G., Everatt, J., Bhiman, J.N., Scheepers, C., et al. (2022). Clinical severity of SARS-CoV-2 Omicron BA.4 and BA.5 lineages compared to BA.1 and Delta in South Africa. Nat Commun 13, 5860.

Xu, E., Xie, Y., and Al-Aly, Z. (2022). Long-term neurologic outcomes of COVID-19. Nat Med 28, 2406–2415.

Yan, K., Dumenil, T., Tang, B., Le, T.T., Bishop, C.R., Suhrbier, A., and Rawle, D.J. (2022). Evolution of ACE2-independent SARS-CoV-2 infection and mouse adaption after passage in cells expressing human and mouse ACE2. Virus Evol 8, veac063.

Yan, K., Rawle, D.J., Le, T.T., and Suhrbier, A. (2021). Simple rapid in vitro screening method for SARS-CoV-2 anti-virals that identifies potential cytomorbidity-associated false positives. Virol J 18, 123.

Yang, A.C., Kern, F., Losada, P.M., Agam, M.R., Maat, C.A., Schmartz, G.P., Fehlmann, T., Stein, J.A., Schaum, N., Lee, D.P., et al. (2021). Dysregulation of brain and choroid plexus cell types in severe COVID-19. Nature 595, 565–571.

Yinda, C.K., Port, J.R., Bushmaker, T., Offei Owusu, I., Purushotham, J.N., Avanzato, V.A., Fischer, R.J., Schulz, J.E., Holbrook, M.G., Hebner, M.J., et al. (2021). K18-hACE2 mice develop respiratory disease resembling severe COVID-19. PLoS Pathog 17, e1009195.

Yu, P., Deng, W., Bao, L., Qu, Y., Xu, Y., Zhao, W., Han, Y., and Qin, C. (2022). Comparative pathology of the nasal epithelium in K18-hACE2 Tg mice, hACE2 Tg mice, and hamsters infected with SARS-CoV-2. Vet Pathol 59, 602–612.

Yue, C., Song, W., Wang, L., Jian, F., Chen, X., Gao, F., Shen, Z., Wang, Y., Wang, X., and Cao, Y. (2023). ACE2 binding and antibody evasion in enhanced transmissibility of XBB.1.5. Lancet Infect Dis 23, 278–280.

Zhang, B.Z., Chu, H., Han, S., Shuai, H., Deng, J., Hu, Y.F., Gong, H.R., Lee, A.C., Zou, Z., Yau, T., et al. (2020). SARS-CoV-2 infects human neural progenitor cells and brain organoids. Cell Res 30, 928–931.

Zhao, H., Lu, L., Peng, Z., Chen, L.-L., Meng, X., Zhang, C., Ip, J.D., Chan, W.-M., Chu, A.W.-H., Chan, K.-H., et al. (2022). SARS-CoV-2 Omicron variant shows less efficient replication and fusion activity when compared with Delta variant in TMPRSS2-expressed cells. Emerging Microbes & Infections 11, 277–283.

Zheng, J., Wong, L.R., Li, K., Verma, A.K., Ortiz, M.E., Wohlford-Lenane, C., Leidinger, M.R., Knudson, C.M., Meyerholz, D.K., McCray, P.B., Jr., et al. (2021). COVID-19 treatments and pathogenesis including anosmia in K18-hACE2 mice. Nature 589, 603–607.

