## Supplementary Figures for "Increased neurovirulence of omicron BA.5 and XBB variants over BA.1 in K18-hACE2 mice and human brain organoids"

**A**

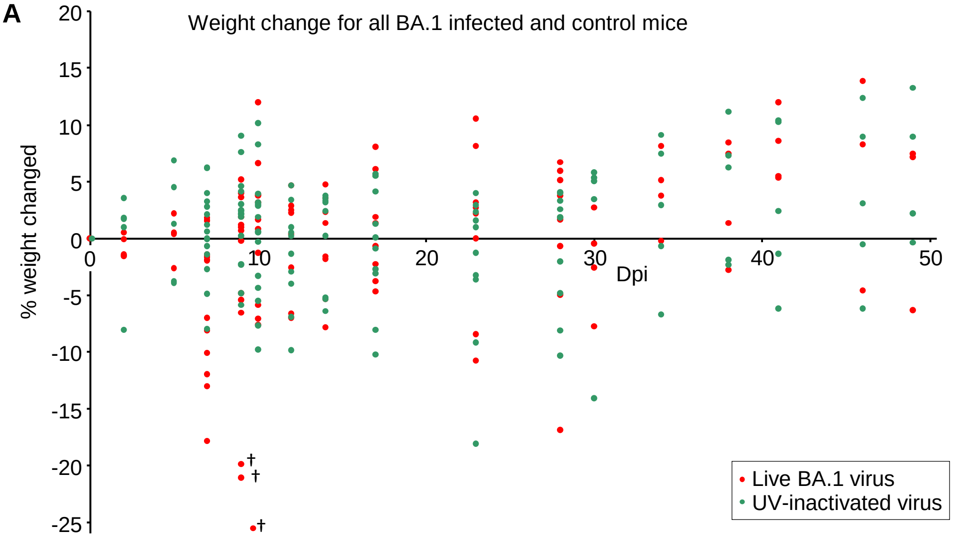


**B**

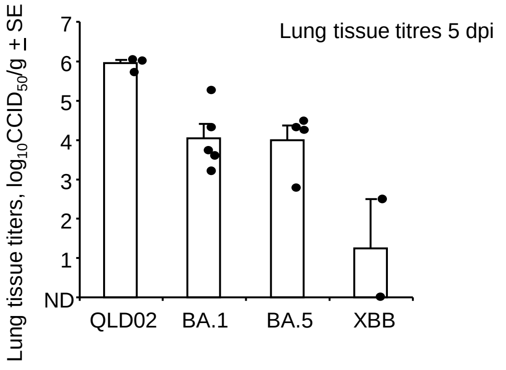


**Supplementary Fig. 1. Detailed weight loss data and lung tissue titres. A** Expanded data for Fig. 1A showing all weight measurements. There were n=24 mice in total infected with BA.1 and n=25 mice inoculated with UV-inactivated BA.1. Some mice were euthanized at specific time points to assess tissue titres (Fig. 1D), others n=3 (†), were euthanized as they reach the ethically defined end point (>20% weight loss). Note that inoculation with UV-inactivated virus (at the same protein dose as live virus) by itself causes weight loss, although usually <15%. **B** Lung tissue titres. All mice were euthanized on 5 dpi. Except for BA.1-infected mice, mice had reached ethically defined end points requiring euthanasia. ND – not detected (limit of detection ≈ 2 $log_{10}CCID_{50}/g$). (Data from 2-3 independent experiments). For the two XXB-infected mice euthanized 6 dpi lung titers where 2.6 $log_{10}CCID_{50}/g$ and ND.

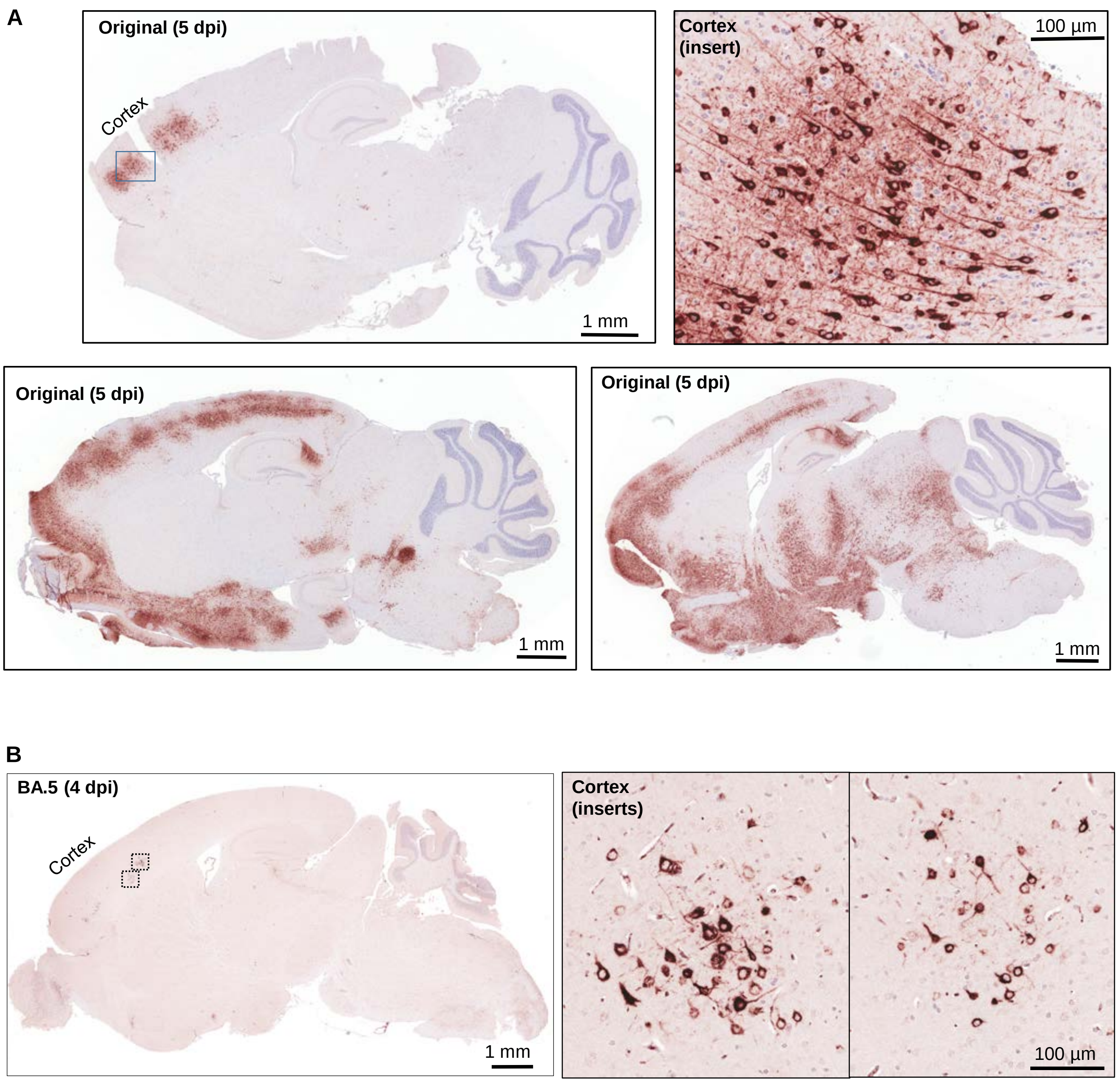


**Supplementary Fig. 2. Anti-spike monoclonal antibody straining of brains by immuno-histochemistry (IHC). A** IHC of brains from K18-hACE2 mice infected with original strain isolate, illustrating the range of IHC staining. Top right shows an enlarged image from the cortex indicating infection of neurons. **B** As for a but infection with omicron BA.5. Despite relatively low level of IHC staining in the brain this mouse reached ethically defined endpoint for weight loss requiring euthanasia by day 4.

**A**

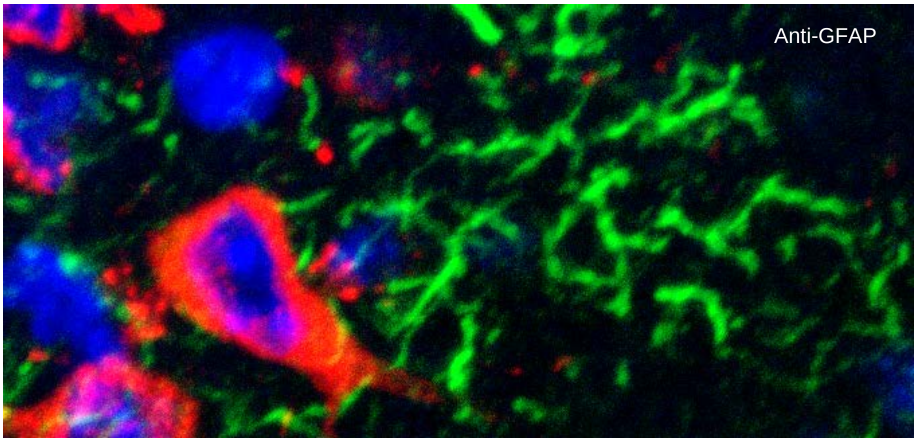


**B**

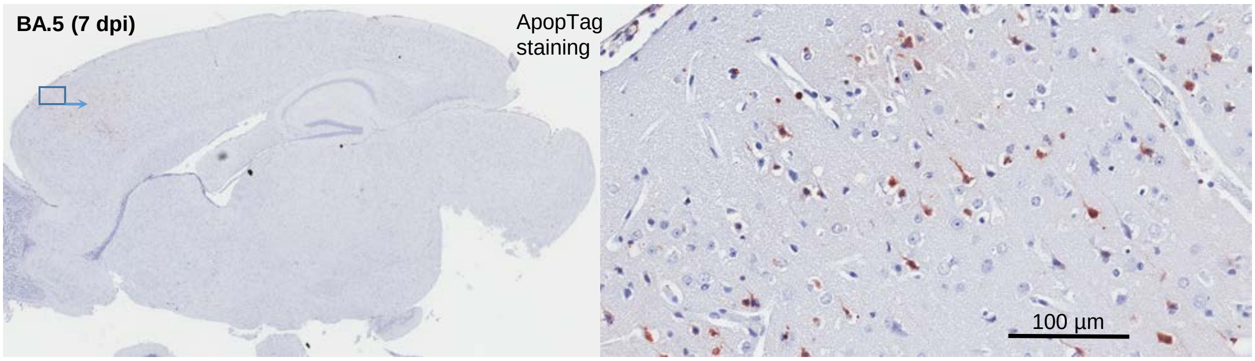


**Supplementary Fig. 3**. **Anti-GFAP and ApopTag staining. A** Image to illustrate anti-GFAP staining is working. Staining with anti-GFAP (as in Fig. 3, Reactive astrocytes) shows classical intermediate filament staining comprised largely of glial fibrillary acidic protein (GFAP) (green), next to a BA.5 infected cell (red). Blue – DAPI (nuclei). **B** ApopTag staining (brown) of the brain shown in Fig. 2B. Insert enlargement shown on the right.

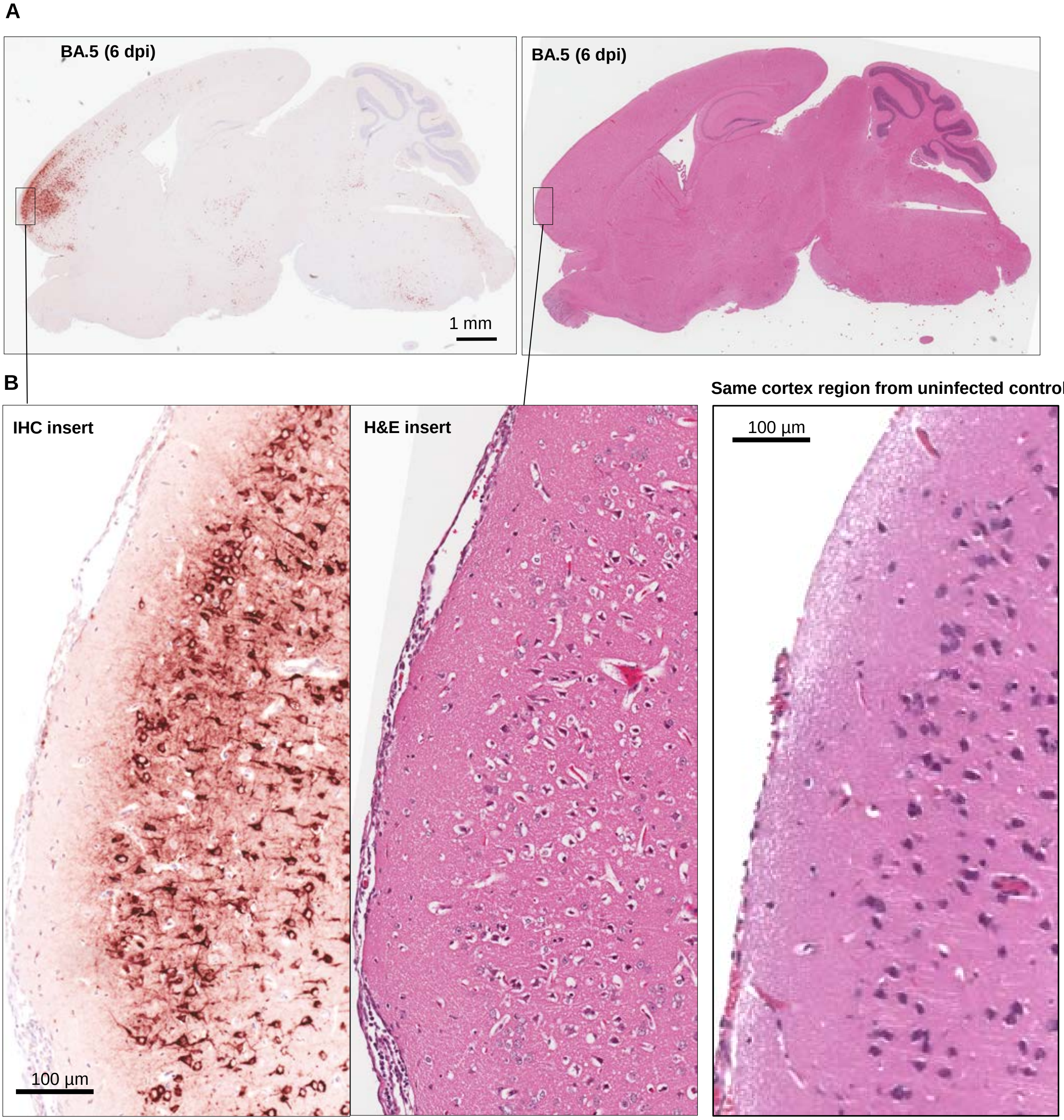


**Supplementary Fig. 4**. **Anti-spike IHC. A** Anti-spike monoclonal antibody straining by immuno-histochemistry (IHC) of brain from a BA.5 infected K18-hACE2 mouse (left) and H&E staining of a section from the same block (right). **B** Inserts showing association of viral antigen detection in the cortex (IHC positive staining) with high density of H&E lesions (primarily vacuolation). A control is shown bottom right; H&E staiing of the same region of the cortex in a control uninfected animal.

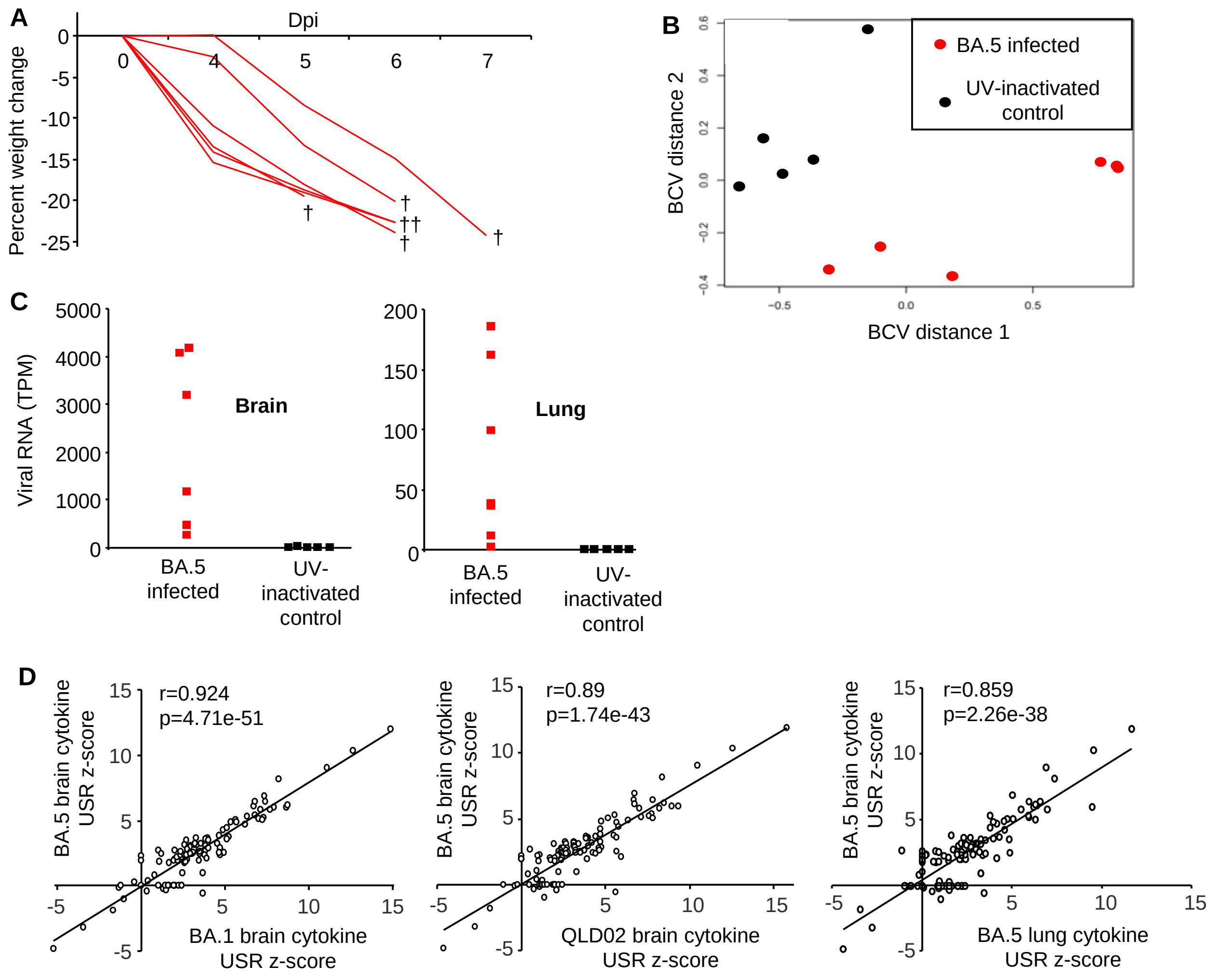


**Supplementary Fig. 5. RNA-Seq of brains from BA.5 infected K18-hACE2 mice.**

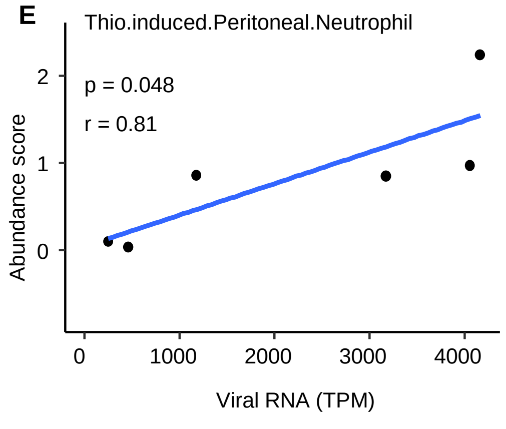


**A** Weight loss for BA.5 infected K18-hACE2 mice used for RNA-Seq (no measurements were taken on 2 or 3 dpi). Half the brain was used for RNA-Seq and the other half was analysed by histology. Mice reached ethically defined end points on 5 dpi (n=1), 6 dpi (n=4), and 7 dpi (n=1), brains were harvested at these time points and used for RNA-Seq. **B** PCA plot showing segregation between brain samples from K18-hACE2 mice that were infected with BA.5 (n=6) or inoculated with UV-inactivated virus (n=5). **C** Viral RNA (TPM) in brain and lung of euthanized mice. **D** Pearson correlations for cytokine IPA USR z-scores between indicated groups. When an USR was absent (not significant) for one group but present (significant) for the other, a z-score of 0 was given to the former. The BA.1 data set was derived from the 3 mice that died 9/10 dpi (n=3), with controls inoculated with UV-inactivated virus (n=5). The QLD02 data set was derived from the 5 mice euthanized 5 dpi (n=5), with brains from naive mice used as controls (n=5). The BA.5 lung data set was obtained from the same mice as for the brains. **E** From Fig. 5B (as for 5C) showing significant Pearson correlation between the indicated neutrophil cell abundance scores and viral RNA levels.

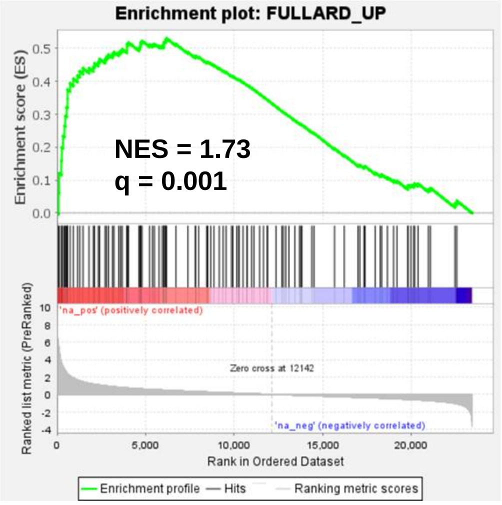


DEGs upregulated in brains of COVID-19 patients (Fullard study) enriched in gene list from brains of BA.5 infected K18-hACE2 mice

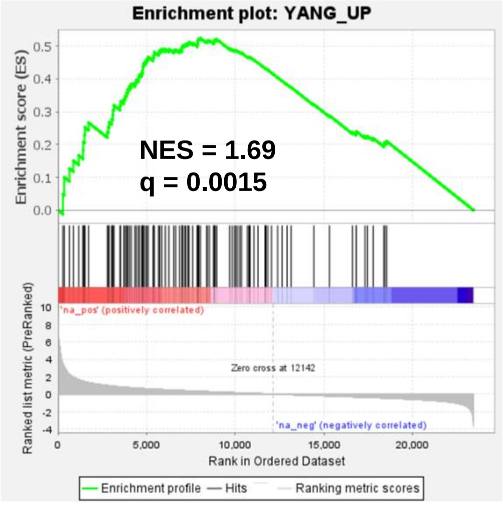


DEGs upregulated in brains of COVID-19 patients (Yang study) enriched in gene list from brains of BA.5 infected K18-hACE2 mice

**Supplementary Fig. 6. GSEAs show significant concordance in gene expression profiles in brains of severe COVID-19 patients and BA.5 infected K18-hACE2 mice.** Gene expression data from two studies on COVID-19 patient brains (Fullard et al., 2021; Yang et al, 2021) contained 20 and 45 gene expression data sets, respectively. Filters were applied to each data set; $q<0.05$ and $\log_2$ fold change >1. All genes that passed these filters were then concatenated into a single DEG list for each study. All human gene names were converted to their mouse orthologue (Bishop et al., 2022). Up and down-regulated DEGs from these lists were then separately used in Gene Set Enrichment Analyses (GSEAs) using the ranked (fold change) gene list from brains of BA.5-infected K18-hACE2 mice. GSEAs for down-regulated DEGs did not reach significance, a feature observed previously (Bishop et al., 2022).

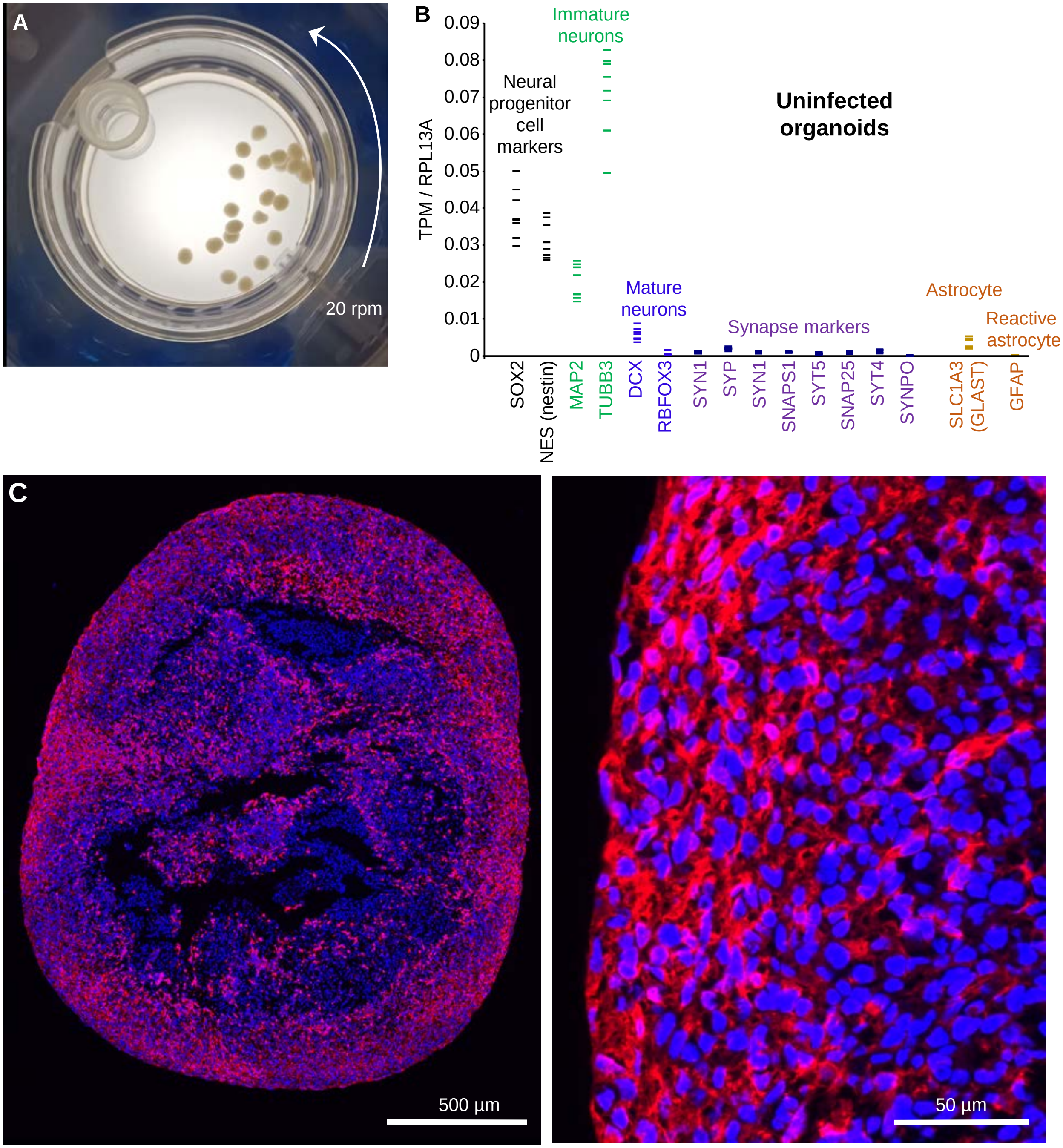


**Supplementary Fig. 7. Uninfected organoids, RNA-Seq and IHC. A** Photograph of "mini-brains" grown in a rotating CelVivo Clinostar incubator. **B** RNA-Seq of uninfected organoids (n=8); expression of markers normalized to the house keeping gene, RPL13A (TPM - transcripts per million) (Supplementary Table 1, TPM). Markers of immature neurons dominated, with such cells often able to retain stem cell marker expression. mRNA expression of markers of mature neurons were lower, with low level expression of synapse genes. Expression of the astrocyte marker, GLAST, was also relatively low suggesting only a small proportion of the cells in the organoid had differentiated into astrocytes. Expression of GFAP, a marker of reactive astrocytes, was very low/undetectable. **C** IHC using anti-MAP2 antibody (Cat# M9942, Sigma) and Opal 650 Reagent Kits (Geneworks) showing widespread staining of cells in the organoid (red). Nuclei were stained with DAPI (blue). For IHC method see Morgan et al., 2022. Images were acquired by Aperio Scanscope FL and compiled in ImageJ (Fiji 1.54b).

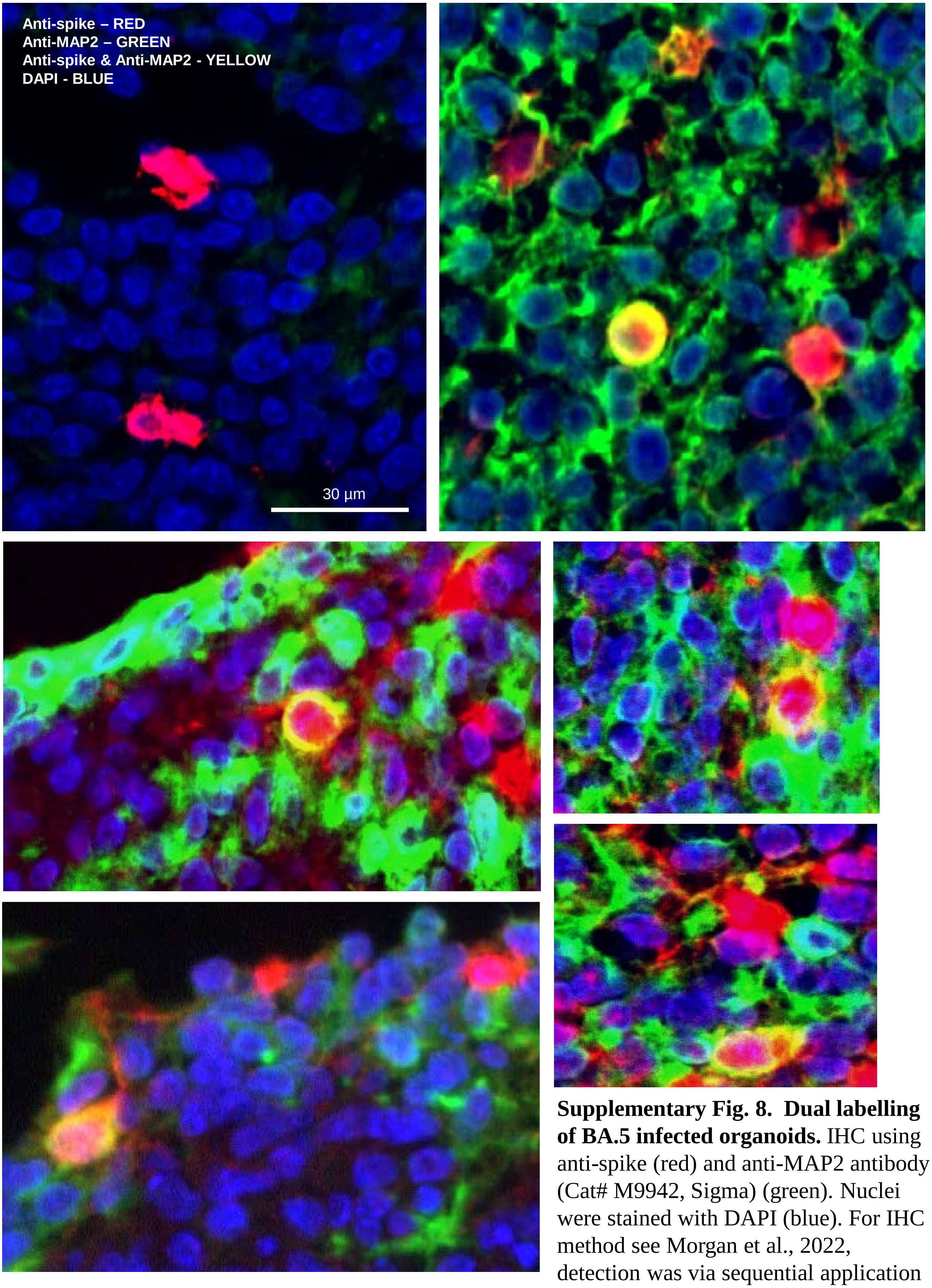


**Supplementary Fig. 8. Dual labelling of BA.5 infected organoids.** IHC using anti-spike (red) and anti-MAP2 antibody (Cat# M9942, Sigma) (green). Nuclei were stained with DAPI (blue). For IHC method see Morgan et al., 2022, detection was via sequential application of Opal 650 and Opal 570 Reagent Kits (Geneworks). Images were acquired by Aperio Scanscope FL and compiled in ImageJ (Fiji 1.54b). Top left shows infection of MAP2 negative cells. Yellow in the remaining images shows infection of MAP2 positive cells (red green overlay).

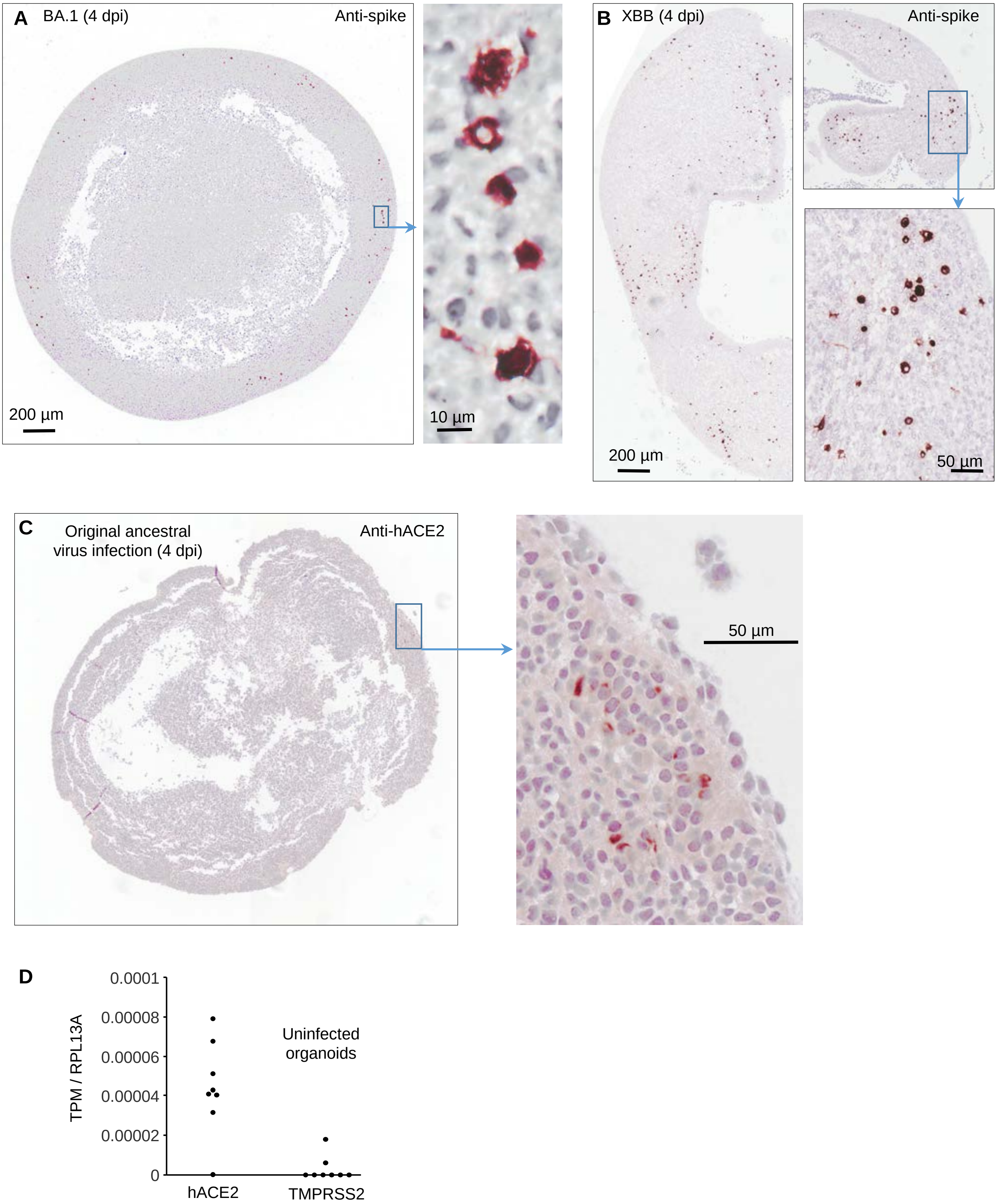


**Supplementary Fig. 9. Human cortical brain organoid infection.** **A** As for Fig 6A but infection with BA.1. **B** As for A but infection with XBB. **C** IHC with anti-hACE2 for the organoid shown in Fig. 6C, corresponding to the region (in a parallel section) of viral infection with the original ancestral virus. **D** RNA-Seq data (as in Supplementary Fig. 7B) showing overall expression of hACE2 mRNA was low, with TMPRSS2 mRNA often undetectable (n=8; Supplementary Table 2, TPM).

**A** SARS-CoV-2$_{QLD02}$ (original ancestral isolate) infected K18-hACE2 mice

**Brains** — Nucleotide sequence — Allele freq.

```
Mouse 1    ATGCGCTAGTTAT                  TCTCCTCGGCGGGCACGTAGTGTAGCTAGT  1.0
Mouse 2    ATGCGCTAGTTAT                  TCTCCTCGGCGGGCACGTAGTGTAGCTAGT  0.99
Mouse 3    ATGCGCTAGTTAT   In frame       TCTCCTCGGCGGGCACGTAGTGTAGCTAGT  0.95
Mouse 4    ATGCGCTAGTTAT   deletion       TCTCCTCGGCGGGCACGTAGTGTAGCTAGT  0.99
Mouse 5    ATGCGCTAGTTAT                  TCTCCTCGGCGGGCACGTAGTGTAGCTAGT  0.17
Mouse 6    ATGCGCTAGTTAT                  TCTCCTCGGCGGGCACGTAGTGTAGCTAGT  0.99
Mouse 7    ATGCGCTAGTTATCAGACTCAGACTAATTCTCCTCGGCGGGCACGTAGTGTAGCTAGT  0
Mouse 8    ATGCGCTAGTTATCAGACTCAGACTAATTCTCCTCGGCGGGCACGTAGTGTAGCTAGT  0
Mouse 9    ATGCGCTAGTTATCAGACTCAGACTAATTCTCCTCGGCGGGCACGTAGTGTAGCTAGT  0
Mouse 10   ATGCGCTAGTTATCAGACTCAGACTAATTCTCCTCGGCGGGCACGTAGTGTAGCTAGT  0
Mouse 11   ATGCGCTAGTTATCAGACTCAGACTAATTCTCCTCGGCGGGCACGTAGTGTAGCTAGT  0
Mouse 12   ATGCGCTAGTTATCAGACTCAGACTAATTCTCCTCGGCGGGCACGTAGTGTAGCTAGT  0
```

**Lungs**

```
Mouse 1    ATGCGCTAGTTATCAGACTCAGACTAATTCTCCTCGGCGGGCACGTAGTGTAGCTAGT  0
Mouse 2    ATGCGCTAGTTATCAGACTCAGACTAATTCTCCTCGGCGGGCACGTAGTGTAGCTAGT  0
Mouse 3    ATGCGCTAGTTATCAGACTCAGACTAATTCTCCTCGGCGGGCACGTAGTGTAGCTAGT  0
Mouse 4    ATGCGCTAGTTATCAGACTCAGACTAATTCTCCTCGGCGGGCACGTAGTGTAGCTAGT  0
Mouse 5    ATGCGCTAGTTATCAGACTCAGACTAATTCTCCTCGGCGGGCACGTAGTGTAGCTAGT  0
Mouse 6    ATGCGCTAGTTATCAGACTCAGACTAATTCTCCTCGGCGGGCACGTAGTGTAGCTAGT  0
Mouse 7    ATGCGCTAGTTATCAGACTCAGACTAATTCTCCTCGGCGGGCACGTAGTGTAGCTAGT  0
Mouse 8    ATGCGCTAGTTATCAGACTCAGACTAATTCTCCTCGGCGGGCACGTAGTGTAGCTAGT  0
Mouse 9    ATGCGCTAGTTATCAGACTCAGACTAATTCTCCTCGGCGGGCACGTAGTGTAGCTAGT  0
Mouse 10   ATGCGCTAGTTATCAGACTCAGACTAATTCTCCTCGGCGGGCACGTAGTGTAGCTAGT  0
Mouse 11   ATGCGCTAGTTATCAGACTCAGACTAATTCTCCTCGGCGGGCACGTAGTGTAGCTAGT  0
Mouse 12   ATGCGCTAGTTATCAGACTCAGACTAATTCTCCTCGGCGGGCACGTAGTGTAGCTAGT  0
                       Q  T  Q  T  N  S  P  R  R  A  R
```

**B** Homologous deletion found in human samples (SRA)

**Deletion** CAGACTCAGACTAAG/T (Q T Q T K/N)

| Bioproject | Sample | Depth | Allele Freq. |
|---|---|---|---|
| PRJNA639791 | SRR12023946 | 11855 | 0.119 |
| PRJNA639791 | SRR12023947 | 6999 | 0.081 |
| PRJNA639791 | SRR12023948 | 5201 | 0.046 |
| PRJNA688321 | SRR13319470 | 44 | 0.023 |
| PRJNA688321 | SRR13319472 | 47 | 0.021 |
| PRJNA688321 | SRR13319496 | 37 | 0.027 |
| PRJNA688321 | SRR13319497 | 38 | 0.026 |
| PRJNA688321 | SRR13319501 | 55 | 0.036 |
| PRJNA688321 | SRR13319502 | 66 | 0.015 |
| PRJNA688321 | SRR13319503 | 74 | 0.027 |
| PRJNA688321 | SRR13319504 | 70 | 0.014 |

**C** SARS-CoV-2 (original ancestral isolate) infected hamsters

**Brain** (with deletion n=5 reads)

```
CAGGTATATGCGCTAGTTATCAGACTCAGCAATCCATCATTGCCTACACTATGTC
CAGGTATATGCGCTAGTTATCAGACTCAGCAATCCATCATTGCCTAC
CAGGTATATGCGCTAGTTATCAGACTCAGCAATCCATCATTGCCTACAC
CAGGTATATGCGCTAGTTATCAGACTCAGCAATCCATCATTGCCTACACTATGTCACTTGGTGCAGAAAATTCAGTTGCTTACA
   GTATATGCGCTAGTTATCAGACTCAGCAATCCATCATTGCCTACACTATGTCACTTGGTGCAGAAAATTCAGTTGCTTACA
```

(without deletion n=1 read) — In frame deletion

```
CAGGTATATGCGCTAGTTATCAGACTCAGACTAATTCTCCTCGGCGGGCACGTAGTGTAGCTAGTCAATCCATCATTGCCTACA
```

**Lung** (without deletion n=42 reads, 14 shown)

```
CAGGTATATGCGCTAGTTATCAGACTCAGACTAATTCTCCTCGGCGGGCACGTAGTGTAGCTAGTCAATCCATCATTGCCTACA
CAGGTATATGCGCTAGTTATCAGACTCAGACTAATTCTCCTCGGCGAGCACGTAGTGTAGCTAGTCAATCCATCATTGCCTACA
CAGGTATATGCGCTAGTTATCAGACTCAGACTAATTCTCCTCGGCGGGCACGTAGTGTAGCTAGTCAATCCATCATTGCCTACA
CAGGTATATGCGCTAGTTATCAGACTCAGACTAATTCTCCTCGGCGGGCACGTAGTGTAGCTAGTCAATCCATCATTGCCTACA
                            GACTAATTCTCCTCGGCGGGCACGTAGTGTAGCTAGTCAATCCATCATTGCCTACA
CAGGTATATGCGCTAGTTATCAGACTCAGACTAATTCTCCTCGGCGGGCACGTAGTGTAGCTAGTCAATCCATCATTGCCTACA
                             ACTAATTCTCCTCGGCGAGCACGTAGTGTAGCTAGTCAATCCATCATTGCCTACA
CAGGTATATGCGCTAGTTATCAGACTCAGACTAATTCTCCTCGGCGGGCACGTAGTGTAGCTAGTCAATCCATCATTGCCTACA
CAGGTATATGCGCTAGTTATCAGACTCAGACTAATTCTCCTCGGCGAGCACGTAGTGTAGCTAGTCAATCCATCATTGCCTACA
CAGGTATATGCGCTAGTTATCAGACTCAGACTAATTCTCCTCGGCGGGCACGTAGTGTAGCTAGTCAATCCATCATTGCCTACA
CAGGTATATGCGCTAGTTATCAGACTCAGACTAATTCTCCTCGGCGAGCACGTAGTGTAGCTAGTCAATCCATCATTGCCTACA
 AGGTATATGCGCTAGTTATCAGACTCAGACTAATTCTCCTCGGCGGGCACGTAGTGTAGCTAGTCAATCCATCATTGCCTACA
  GGTATATGCGCTAGTTATCAGACTCAGACTAATTCTCCTCGGCGGGCACGTAGTGTAGCTAGTCAATCCATCATTGCCTACA
    TATATGCGCTAGTTATCAGACTCAGACTAATTCTCCTCGGCGGGCACGTAGTGTAGCTAGTCAATCCATCATTGCCTACA
                    Q  T  Q  T  N  S  P  R  R  A  R  S  V  A  S  Q
```

Polybasic (R R A R) — Furin

**D** Homologous deletions found in human samples (SRA)

**Deletion** ACT-AAG/T-TCTCATCGGCGGGCACGTAGTGTAGCTAGT (T K/N S H R R A R S V A S)

| Bioproject | Sample | Depth | Allele Freq. |
|---|---|---|---|
| PRJNA639791 | SRR12023946 | 11836 | 0.0048 |
| PRJNA639791 | SRR12023947 | 7021 | 0.0051 |
| PRJNA639791 | SRR12023948 | 5221 | 0.0048 |

**Deletion** T-AAG/T-TCTCATCGGCGGGCA-CG (K/N S H R R A R)

| Bioproject | Sample | Depth | Allele Freq. |
|---|---|---|---|
| PRJNA639791 | SRR12023946 | 11886 | 0.005 |
| PRJNA639791 | SRR12023947 | 7046 | 0.006 |
| PRJNA639791 | SRR12023948 | 5236 | 0.007 |
| PRJNA688321 | SRR13319472 | 28 | 0.036 |
| PRJNA688321 | SRR13319473 | 26 | 0.038 |
| PRJNA688321 | SRR13319498 | 42 | 0.048 |
| PRJNA688321 | SRR13319499 | 38 | 0.026 |
| PRJNA688321 | SRR13319504 | 43 | 0.023 |
| PRJNA786802 | SRR17155278 | 691 | 0.001 |
| PRJNA786802 | SRR17155279 | 336 | 0.003 |
| PRJNA786802 | SRR17155280 | 652 | 0.008 |

**Supplementary Fig. 10. Loss of functional furin cleavage site. A** QTQTN flanking the furin cleavage site, is often lost in a large proportion of virus from brains of K18-hACE2 mice infected with SARS-CoV-2$_{QLD02}$. This loss was not seen virus from lungs of the same mice (SRA Bioproject PRJNA813692). Red and green indicate the codons flanking the deletion sites. Purple indicates the deleted sequences. QTQTN is required for a functional furin cleavage site (Vu *et al.*, 2022. PNAS 119; e2205690119. **B** The same or homologous deletion also exists in human samples (submitted to the NCBI SRA). **C** RNA-Seq data from brains and lungs of hamsters infected with SARS-CoV-2 original strain (WA1) (SRA Bioproject PRJNA837993) (Frere *et al*. 2022. Sci Trans Med 14; eabq3059). Of the 6 sequence reads from brain spanning the furin clevage site, 5 had deletions (TNSPRRARSVAS). This deletion was not identified in any lung sequences. **D** The same, homologous and related deletions can also be found in human samples.

METHOD: To identify RNA-Seq datasets of SARS-CoV-2-infected humans, metadata from the Sequence Read Archive (SRA) was searched using Google BigQuery and the NCBI Taxonomy Analysis Tool. To search for nucleotide deletions in SARS-CoV-2 infected mouse and human RNA-Seq data, fastq files were aligned to the SARS-CoV-2 genome using Bowtie2 v2.5.0. Alignments were pre-processed using Lofreq v2.1.3.1, Genome Analysis Toolkit v4.2.4.1, and Samtools v1.6. Deletions were called using Lofreq with a minimum p-value threshold of 0.05.

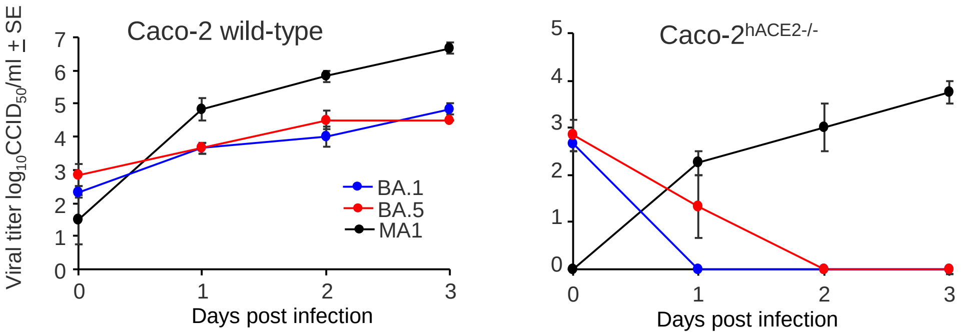


**Supplementary Fig. 11. No replication of BA.5 in hACE2 negative cell lines.** Caco-2 cells express hACE2 and can be infected by BA.1 and BA.5, with a mouse adapted virus, MA1, also able to infect these cells. MA1 is able also to productively infect Caco-2 cells wherein hACE2 has been deleted by CRISPR (Yan *et al*, 2022). This ability to infect hACE2-negative cells was not seen for BA.1 or BA.5.

```
BA.1   1     MFVFLVLLPLVSSQCVNLTTRTQLPPAYTNSFTRGVYYPDKVFRSSVLHSTQDLFLPFFS  60
BA.5   1     ..................I....---S.................................  57

61     NVTWFHVISGTNGTKRFDNPVLPFNDGVYFASIEKSNIIRGWIFGTTLDSKTQSLLIVNN  120
58     ......A.........................T...........................  117

121    ATNVVIKVCEFQFCNDPFLD---HKNNKSWMESEFRVYSSANNCTFEYVSQPFLMDLEGK  177
118    ....................VYY.....................................  177

178    QGNFKNLREFVFKNIDGYFKIYSKHTPISEPEDLPQGFSALEPLVDLPIGINITRFQTLL  237
178    ............................NLGR............................  237

238    ALHRSYLTPGDSSSGWTAGAAAYYVGYLQPRTFLLKYNENGTITDAVDCALDPLSETKCT  297
238    ............................................................  297

298    LKSFTVEKGIYQTSNFRVQPTESIVRFPNITNLCPFDEVFNATRFASVYAWNRKRISNCV  357
298    ............................................................  357

358    ADYSVLYNLAPFFTFKCYGVSPTKLNDLCFTNVYADSFVIRGDEVRQIAPGQTGNIADYN  417
358    ........F....A............................N..S..............  417

418    YKLPDDFTGCVIAWNSNKLDSKVSGNYNYLYRLFRKSNLKPFERDISTEIYQAGNKPCNG  477   RBD
418    .......................G.....R..............................  477

478    VAGFNCYFPLRSYSFRPTYGVGHQPYRVVVLSFELLHAPATVCGPKKSTNLVKNKCVNFN  537
478    ...V......Q..G..............................................  537

538    FNGLKGTGVLTESNKKFLPFQQFGRDIADTTDAVRDPQTLEILDITPCSFGGVSVITPGT  597
538    ....T.......................................................  597

598    NTSNQVAVLYQGVNCTEVPVAIHADQLTPTWRVYSTGSNVFQTRAGCLIGAEYVNNSYEC  657
598    ............................................................  657

658    DIPIGAGICASYQTQTKSHRRARSVASQSIIAYTMSLGAENSVAYSNNSIAIPTNFTISV  717
658    ............................................................  717

718    TTEILPVSMTKTSVDCTMYICGDSTECSNLLLQYGSFCTQLKRALTGIAVEQDKNTQEVF  777
718    ............................................................  777

778    AQVKQIYKTPPIKYFGGFNFSQILPDPSKPSKRSFIEDLLFNKVTLADAGFIKQYGDCLG  837
778    ............................................................  837

838    DIAARDLICAQKFKGLTVLPPLLTDEMIAQYTSALLAGTITSGWTFGAGAALQIPFAMQM  897
838    .............N..............................................  897

898    AYRFNGIGVTQNVLYENQKLIANQFNSAIGKIQDSLSSTASALGKLQDVVNHNAQALNTL  957
898    ............................................................  957

958    VKQLSSKFGAISSVLNDIFSRLDKVEAEVQIDRLITGRLQSLQTYVTQQLIRAAEIRASA  1017
958    ..................L.........................................  1017

1018   NLAATKMSECVLGQSKRVDFCGKGYHLMSFPQSAPHGVVFLHVTYVPAQEKNFTTAPAIC  1077
1018   ............................................................  1077

1078   HDGKAHFPREGVFVSNGTHWFVTQRNFYEPQIITTDNTFVSGNCDVVIGIVNNTVYDPLQ  1137
1078   ............................................................  1137

1138   PELDSFKEELDKYFKNHTSPDVDLGDISGINASVVNIQKEIDRLNEVAKNLNESLIDLQE  1197
1138   ............................................................  1197

1198   LGKYEQYIKWPWYIWLGFIAGLIAIVMVTIMLCCMTSCCSCLKGCCSCGSCCKFDEDDSE  1257
1198   ............................................................  1257

1258   PVLKGVKLHYT  1268
1258   ...........  1268
```

**Supplementary Fig. 12. BA.1 and BA.5 spike protein sequences**. Spike protein differences between the BA.1 (QIMR01) and BA.5 (QIMR03) isolates used herein. Purple - receptor binding domain. Green – the COR-22 BA.5 isolate (Uraki et al. 2022) has a I in this position. BA.5 (QIMR03) also has a number of other changes from COR-22; NS3 (Orf3a) G49C, NS3 (Orf3a) V48F, NSP3 A386T, NSP2 Q376K, NSP2 I273V, NSP5 L252P, NSP6 L260F, NSP12 T591I, NSP13 M233I, N E136D.
